# Protein Coronas Derived from Cerebrospinal Fluid Enhance the Interactions Between Nanoparticles and Brain Cells

**DOI:** 10.1101/2024.05.31.596763

**Authors:** Nabila Morshed, Claire Rennie, Matthew Faria, Lyndsey Collins-Praino, Andrew Care

## Abstract

Neuronanomedicine harnesses nanoparticle technology for the treatment of neurological disorders. An unavoidable consequence of nanoparticle delivery to biological systems is the formation of a protein corona on the nanoparticle surface. Despite the well-established influence of the protein corona on nanoparticle behavior and fate, as well as FDA approval of neuro-targeted nanotherapeutics, the effect of a physiologically relevant protein corona on nanoparticle-brain cell interactions is insufficiently explored. Indeed, less than 1% of protein corona studies have investigated protein coronas formed in cerebrospinal fluid (CSF), the fluid surrounding the brain. Herein, we utilize two clinically relevant polymeric nanoparticles (PLGA and PLGA-PEG) to evaluate the formation of serum and CSF protein coronas. LC-MS analysis revealed distinct protein compositions, with selective enrichment/depletion profiles. Following incubation with brain cells, serum and CSF coronas on PLGA particles showed enhanced associations with all cell types as compared to their corresponding corona on PLGA-PEG particles. CSF-derived protein coronas on PLGA nanoparticles, specifically, showed the greatest nanoparticle-cell interactions, with Pearson’s correlation analysis revealing that proteins associated with enhanced nanoparticle-cell interactions were exclusively enriched in this protein corona. This study demonstrates the importance of correct choice of physiologically relevant biological fluids, and its influence on the formation of the protein corona, subsequent nanoparticle-cell interactions.

## 1. Introduction

Neuronanomedicine aims to engineer nanoparticle-based drug delivery systems (NDDSs) for the targeted treatment of brain-related diseases.^[1]^ NDDSs can enhance therapeutic efficacy while reducing side effects by improving drug stability, safety, bioavailability, and mediating drug transport across the blood-brain barrier (BBB) to specific target sites in the brain.^[1]^ Despite such promise, however, the translation of neuronanomedicines into the clinic has been met with limited success. This can be attributed to a poor understanding of how nanoparticles interact with complex biological systems, including the brain.^[2–3]^

Spontaneous coating of nanoparticles with proteins occurs upon contact with biological fluids, forming an outer layer known as the ‘protein corona’. This phenomenon alters the biophysical properties of nanoparticles (i.e., size, shape, dispersity, surface functionality) and is thought to determine both their *in vivo* behavior and fate (i.e., stability, pharmacokinetics, biodistribution, targeting, nanotoxicity) and, in the case of NDDSs, therapeutic performance.^[2, 4]^

Protein coronas are dynamic, with their composition strongly influenced by the physicochemical properties of the nanoparticles, as well as the particular biological fluid to which they are exposed.^[5]^ At present, *in vitro* evaluations of NDDSs typically involve the use of immortalized cell lines in fetal bovine serum (FBS)-supplemented media.^[6–7]^ However, this experimental design overlooks the fact that different biological fluids result in distinct coronas, which can potentially influence their therapeutic efficacy and side effect profile.^[5]^ Indeed, serum/plasma sourced from different species, or even individuals of the same species, form protein coronas that vary in both composition, in turn affecting nanoparticle-cell interactions (e.g., cell uptake, immune responses).^[8]^ Such observations therefore pose a challenge to the effective translation of *in vitro* findings to downstream animal studies and human clinical trials, and highlight the critical necessity for further work in this area.

Brain-targeting NDDSs are designed to enter the brain parenchyma, where they interact directly with circulating cerebrospinal fluid (CSF). This fluid has multiple functions within the brain, including mechanical protection, facilitation of central nervous system (CNS) communication, and the maintenance of biochemical homeostasis through nutrient delivery and waste removal.^[9–10]^ Nevertheless, while CSF composition is well-established (i.e., water, proteins, ions, neurotransmitters, glucose)^[11]^ the adsorption of CSF biomolecules onto the surface of nanoparticles remains understudied. In fact, a systematic review of 470 studies on protein corona formation showed that, among 1826 biological fluid sources, serum (51%), plasma (29%) and cell culture medium (11%) were the most predominantly studied, while CSF accounted for less than 1%.^[12]^

Significantly, in a recent report, Cox *et al*. showed that nanoparticle coronas are likely to undergo significant compositional changes upon traversing the BBB.^[7]^ In a transwell BBB model, ultrasmall gold nanoparticles were introduced into FBS-containing medium ("blood"), before migrating through a layer of brain endothelial cells ("BBB"), into FBS-free medium ("brain"). Of the Top-20 most abundant proteins identified in FBS-derived (“blood”) coronas formed on the nanoparticles, only nine were retained or became enriched after crossing the *in vitro* BBB into the FBS-free medium (“brain”). The other eleven proteins were notably depleted, indicating possible corona shedding and/or exchange during BBB passage. Although this study did not investigate post-BBB interactions between nanoparticles and CSF, it highlights that important differences may exist between the protein corona observed peripherally vs centrally in the nervous system and underscores a significant knowledge gap in current understanding of protein corona behavior in brain-relevant fluids/environments, which may play a role in predicting neuronanomedicine performance.

Motivated by this apparent lack of information, we aimed to investigate how nanoparticles interact with CSF, and how this, in turn, impacts their interactions with brain cells. In comparison to serum (a suitable benchmark fluid), polymeric nanoparticles exposed to CSF were found to develop protein coronas with distinct biophysical characteristics and compositional profiles. Additionally, evaluating the biological effects of corona-nanoparticle complexes on various brain cell types revealed the capacity of specific CSF proteins to augment nanoparticle uptake, where the presence of a CSF corona significantly increased the uptake of PLGA nanoparticles across the representative brain cell subtypes, as compared to bare or even serum coated nanoparticles. Our study therefore sheds light on the role of protein coronas in neuronanomedicine, providing critical insights for refining experimental approaches, and consequently, improving clinical translation.

## 2. Results and discussion

Up until now, CSF-derived coronas have been reported for only a handful of nanoparticles (i.e., gold, silica, polystyrene, carbon), with studies focusing on protein corona composition, while overlooking subsequent influence on nanoparticle behavior (**Supplementary Table S1**).^[13–16]^ In this study (**Scheme 1**), polymeric nanoparticles were incubated with CSF (or serum) to generate corona-nanoparticle complexes. These complexes were characterized *via* proteomic analysis and then incubated with a panel of brain-related cells (i.e. differentiated neurons; microglia; astrocytes), with cellular association, cell viability, and immune responses evaluated. For practical reasons related to CSF volume yield, we elected to use both normal serum and pooled CSF obtained from Sprague-Dawley rats, ensuring species consistency between the two fluids.

We also employed polymeric PLGA and PLGA-PEG nanoparticles as representative brain-targeting NDDSs.^[17]^ These FDA-approved particles are non-toxic, biodegradable, and can be engineered for controlled drug loading/release and passive/active delivery across the BBB into the brain.^[18–19]^ PLGA nanoparticles are often PEGylated to reduce protein corona formation, thereby improving their safety and therapeutic efficacy by mitigating unwanted recognition and interactions inside the body.^[20–22]^ Furthermore, PLGA and PLGA-PEG particles have both been used extensively to study protein corona formation, including that sourced from rats,^[12]^ making them a suitable internal benchmark for this work.

**Scheme 1:**
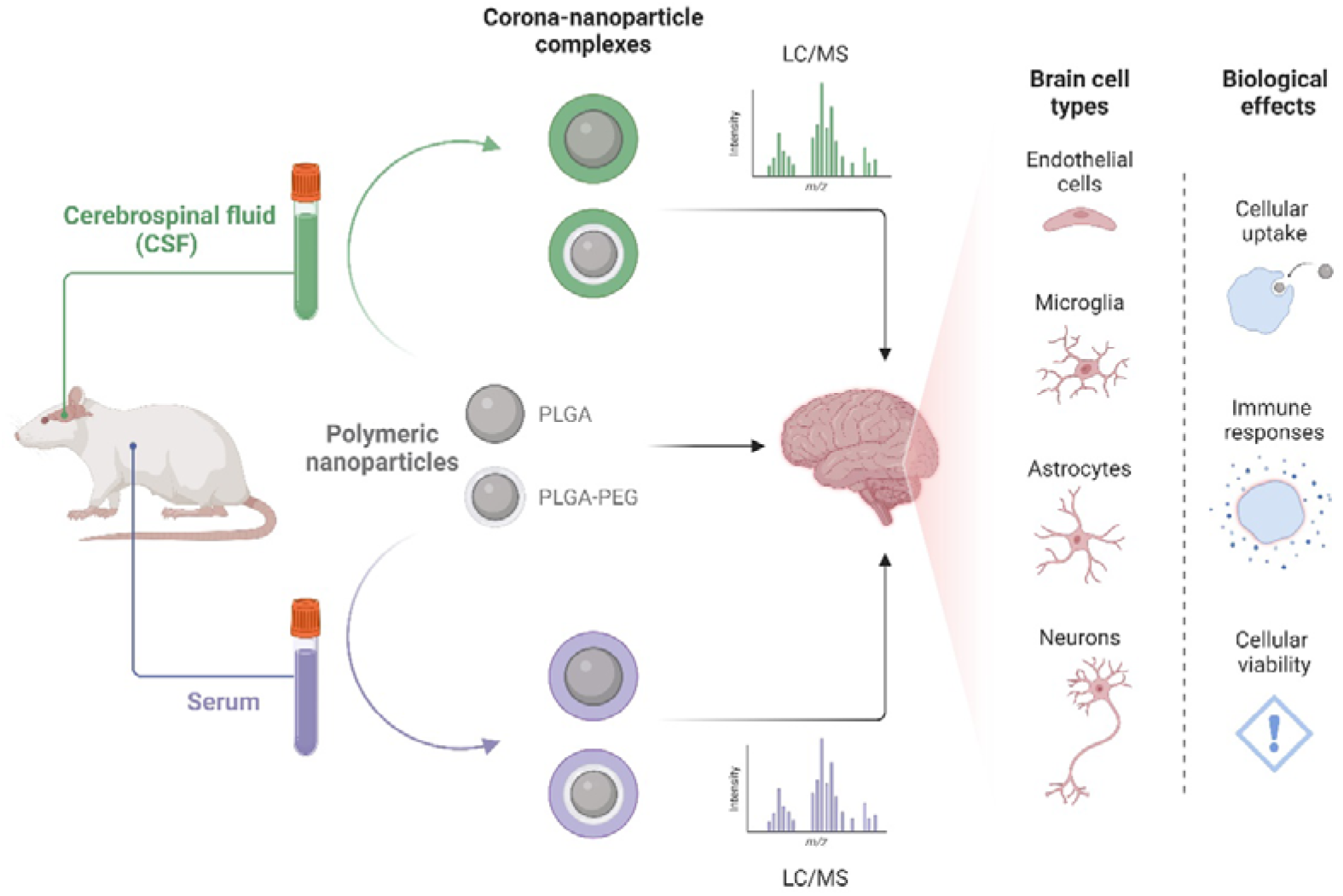
Experimental set-up to study CSF (and serum) corona formation on nanoparticles and their impact on interactions with brain cells.

### Polymeric nanoparticles form protein coronas in CSF

Both PLGA and PLGA-PEG nanoparticles were synthesized using a double emulsion solvent evaporation method ^[23]^ and then characterized. Dynamic light scattering (DLS) showed that PLGA particles had a mean hydrodynamic diameter of 123.7 nm, with a low polydispersity index (PDI = 0.10), indicating uniform size distribution. Zeta potential analysis further determined an overall negative surface charge of -8.1 mV due to exposed carboxyl groups. The PLGA-PEG particles were smaller at 72 nm (PDI = 0.10), possibly due to tighter copolymer packing during synthesis; and exhibited a more negative charge of -33.9 mV that can be attributed to the high negativity of its copolymer. Scanning electron microscopy (SEM) imaging confirmed that each of the polymeric nanoparticles were uniform in their size and spherical shape (**Figure 1 a, g**). Both these nanoparticles were below 200 nm, a size range chosen as <200 nm particles are known to be able to cross the BBB, and can therefore be used as potential neuronanomedicines.^[24]^

**Figure 1:**
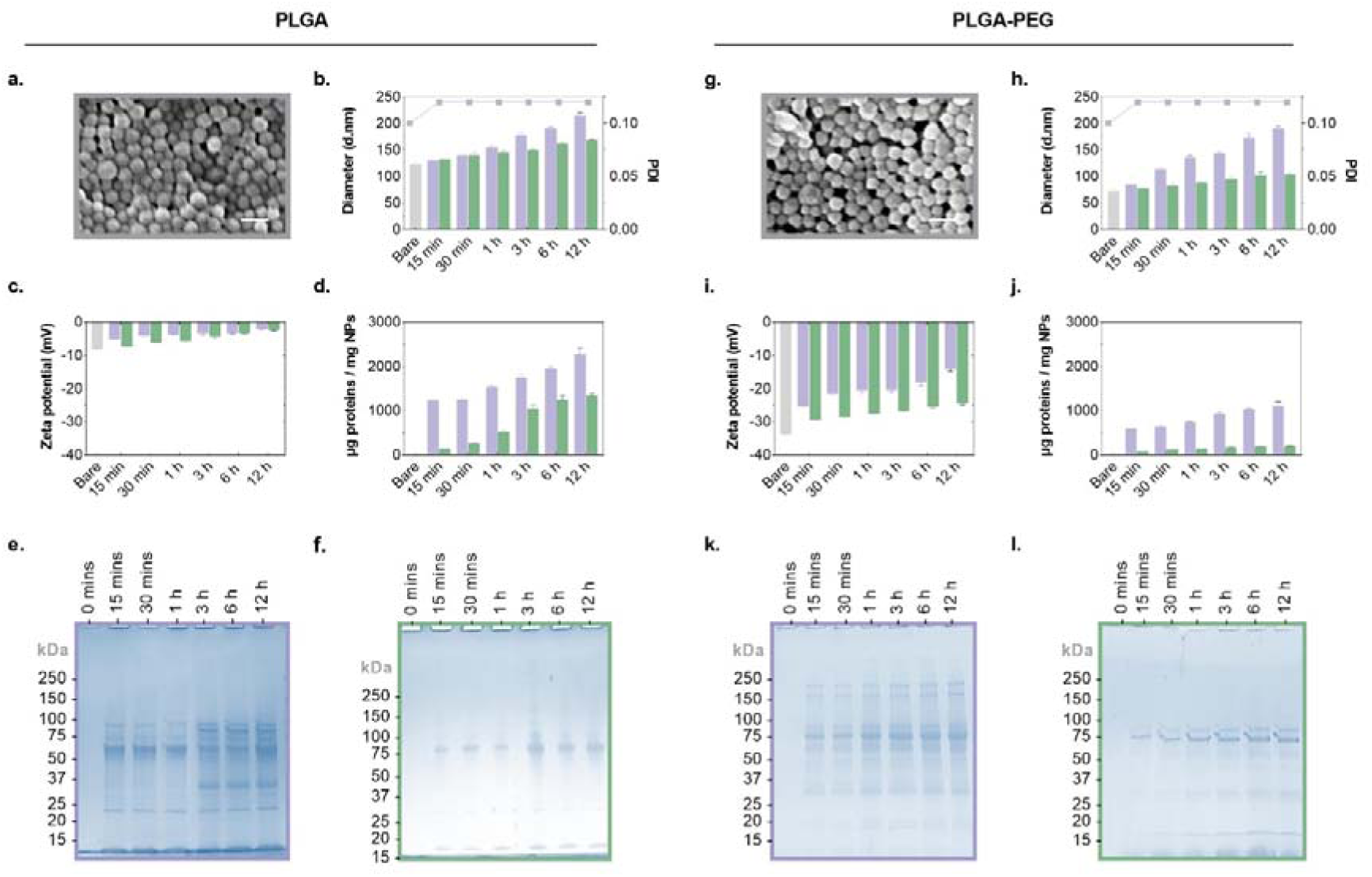
Formation of CSF- and serum-derived protein coronas on polymeric nanoparticles. PLGA **(left panel)** or PLGA-PEG **(right panel)** particles were first visually confirmed for structural formation *via* SEM **(a, g)**. Particles were then incubated with either rat serum (purple) or rat CSF (green) for different periods of time (0, 15 or 30 min, 1, 3, 6, or 12 h), and the resulting corona-nanoparticle complexes subjected to biophysical characterization. Over 12 hours **(b, h)** DLS measurements showed enlargements in particle size (d.nm), but no significant changes in monodispersity (PDI); **(c, i)** Zeta potential analysis (mV) revealed shifts from negative to more positive surface charges; **(d, j)** Total protein mass adsorbed per milligram of nanoparticle (μg/mg) increased; Coomassie stained SDS-PAGE visualized differences in the composition and concentration of protein coronas derived from **(e, k)** serum or **(f, l)** CSF. (Scale bars = 200 μm). Grey = Polymeric nanoparticle; Purple = Serum; Green = CSF. Results presented as mean ± SD.

With these two distinct polymeric nanoparticles in hand, we proceeded with evaluating protein corona formation on their surfaces upon exposure to serum or CSF. After normalizing particle concentrations by surface area, they were individually incubated with either serum or CSF for different time periods (0, 15 or 30 min, 1, 3, 6, or 12 h). Any weakly associated proteins were removed from the particles by washing, leaving behind hard corona coatings.^[25]^ These corona-nanoparticle complexes were then subjected to biophysical characterization (**Figure 1**).

Over 12 h, the diameter of PLGA nanoparticles gradually increased from 124 nm to 215 nm in serum and to 169 nm in CSF (**Figure 1b**), while their surface charge simultaneously shifted from -8 mV to -2.1 mV in serum and -2.4 mV in CSF (**Figure 1c**). PLGA-PEG particles exhibited similar trends, increasing from 72 nm to 190 nm in serum and to 105 nm in CSF (**Figure 1h**); with their surface charge shifting from - 33.9 mV to -14.3 mV in serum and -24.6 mV in CSF **(Figure 1i)**. Notably, all corona-nanoparticle complexes retained a high degree of monodispersity (PDI = 0.12).

In order to quantify the adsorption of fluid proteins onto polymeric particles, the total protein mass absorbed per milligram of nanoparticle (μg/mg) was measured *via* Bradford assay. By the 12 h mark, PLGA particles had steadily adsorbed more proteins from serum (2275 μg/mg) than from CSF (1355 μg/mg) (**Figure 1d**). This ∼50% disparity between the two fluids is likely because serum is a protein-rich biological fluid (60-80 mg/mL), whereas CSF has considerably lower protein levels that range between 0.2-0.5% of those found in blood.^[26]^ When compared to PLGA particles, the PLGA-PEG particles demonstrated substantially lower protein adsorption, with a 2-fold decrease in serum (1110 μg/mg) and a 7-fold decrease in CSF (207 μg/mg) proteins adsorbed (**Figure 1j**). These low-density coronas, especially those derived from CSF, can be attributed to PEG’s ability to repel protein adsorption.^[21]^

We next qualitatively visualized coronas by desorbing proteins from the polymeric nanoparticles and subjecting them to sodium dodecyl-sulfate polyacrylamide gel electrophoresis (SDS-PAGE). Serum and CSF coronas on PLGA nanoparticles displayed distinct protein band patterns and intensities, reflecting their unique compositions and varying concentrations (**Figure 1e, f**). As expected, similar compositions, but significantly lower concentrations, were observed for serum- and CSF-derived coronas on PLGA-PEG particles (**Figure 1k, l**).

Given that the rate of protein exchange would likely have reached equilibrium,^[27–28]^ and that PLGA nanoparticles have previously been detected in circulation at 6 h post injection,^[29–30]^ we elected to use 6 h incubations for the remainder of this study.

### The composition of CSF- and serum-derived protein coronas are distinct

We next set out to profile the protein composition of the serum- and CSF-derived coronas *via* quantitative label-free liquid chromatography-tandem mass spectrometry (LC-MS) analysis. The proteins identified in the biological fluids and adsorbed onto polymeric nanoparticles were categorized based on their molecular weight (MW), hydrophilicity, charge, amino acid content and physiological function (**Figure 2**).

**Figure 2:**
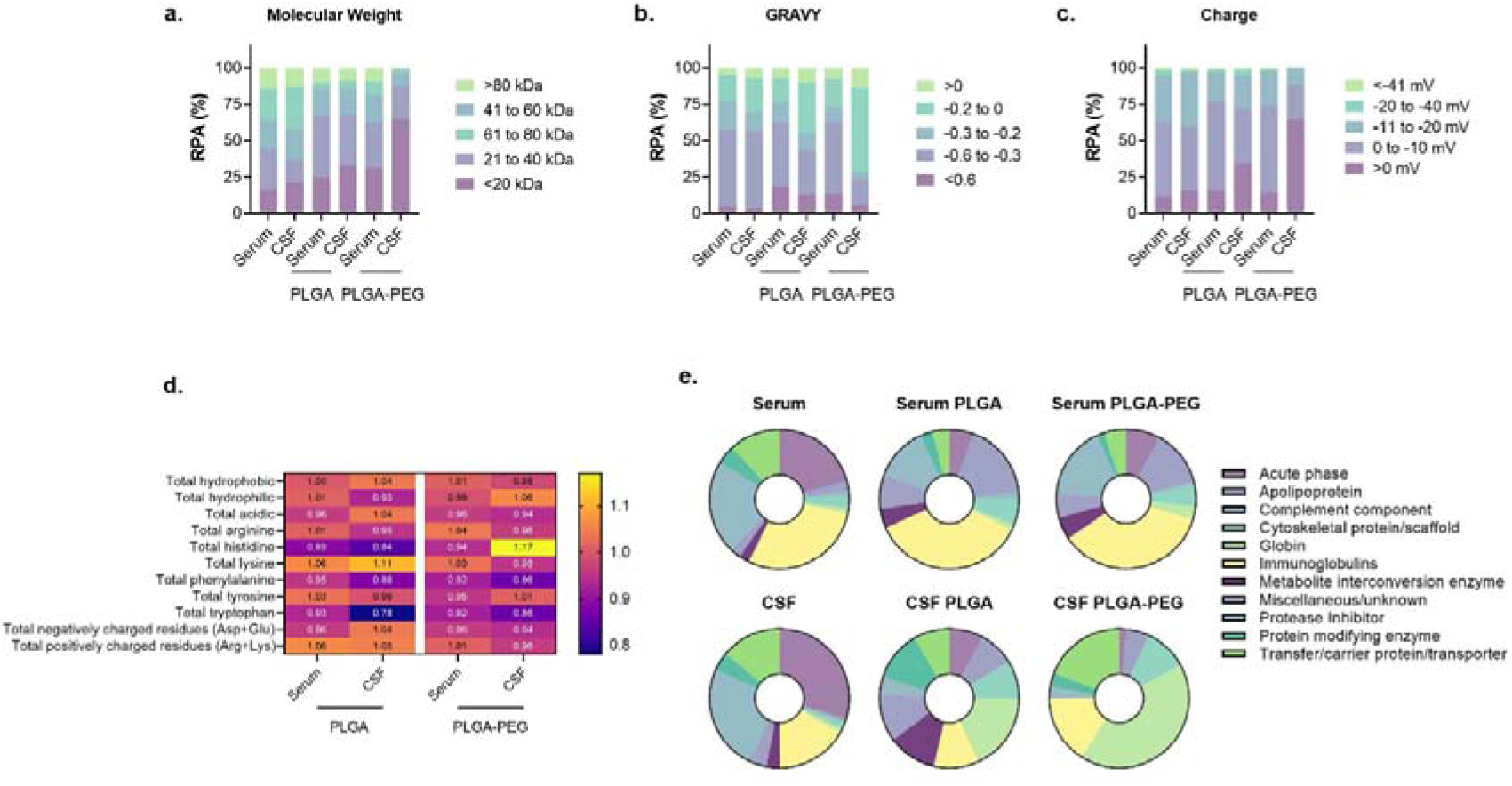
Proteomic analysis of protein coronas formed on polymeric nanoparticles after exposure to serum or CSF. Proteins identified (Top-200) in native biological fluids and their coronas indicated compositional differences in, specifically: **(A)** Molecular weight; **(B)** Hydrophilicity (i.e., GRAVY score); **(C)** Charge; **(D)** Amino acid content and molecular attributes represented by fold-change relative to the respective source fluid (Purple = Reduced; Pink = No change; Yellow = Increased); and **(E)** Physiological function.

When compared to their source fluids, all particle coronas showed greater proportions of low-molecular-weight proteins, with > 60% below 40 kDa (**Figure 2a**). Grand average values of hydropathicity (GRAVY) scores revealed coronas were predominately composed of hydrophilic proteins; however, CSF coronas did exhibit a subtle shift towards more hydrophobic proteins (**Figure 2b**). At physiological pH (7.4), coronas formed were enriched with proteins possessing positive (or more positive) net charges, a characteristic that was particularly pronounced in CSF-derived coatings (**Figure 2c**). A deeper analysis of amino acid composition (**Figure 2d**) indicated that serum coronas closely resembled the native fluid compositions, while CSF coronas on PLGA and PLGA-PEG particles were enriched in positively charged amino acids lysine and histidine, respectively. These findings suggest that CSF corona formation is largely influenced by electrostatic binding interactions between positive proteins and the negatively charged nanoparticle surfaces. In contrast, CSF coronas exhibited a depletion in aromatic residues tryptophan and phenylalanine, which is expected given that these amino acids are instead enriched in coronas on nanoparticles capable of π–π binding interactions, like carbon nanotubes.^[14]^

As depicted in **Figure 2e** (functional classification of all identified Top-200 proteins: **Supplementary Table S2**), the major functional classes in native serum and CSF included acute phase proteins (e.g., albumin), immunoglobulins, protease inhibitors and transfer/carrier proteins/transporters. Serum comprised a higher percentage of immunoglobulins (29%) compared to CSF (17%), while CSF had a greater abundance of acute phase proteins (30%) than serum (21%). In relation to their source fluids, all corona-nanoparticle complexes had lower levels of acute phase proteins, but increased apolipoprotein content. Separately, serum coronas accumulated more immunoglobulins, while CSF coronas instead accrued globins. We further observed that the protein composition of serum coronas were relatively homogeneous across both polymeric nanoparticles, whereas CSF coatings had more compositional diversity. For instance, CSF-derived coronas on PLGA particles showed increases in enzymes related to metabolite interconversion and protein modification, alongside decreases in transfer/carrier proteins/transporters; while CSF coatings on PLGA-PEG particles exhibited opposite trends for these functional classes. Overall, the apparent consistency of serum coronas implies nonspecific protein adsorption onto nanoparticle surfaces, while the diversity seen in CSF coronas suggests some selective protein binding.

Specific proteins exhibit enrichment/depletion in corona-nanoparticle complexes

Our LC-MS analysis revealed the Top-20 most abundant proteins in serum and CSF (**Supplementary Table S3**) and their respective corona-nanoparticle complexes, which are summarized in Table 1 (PLGA) and Table 2 (PLGA-PEG) showing the relative abundance within the Top-20 (RPA). Moreover, the enrichment/depletion of these proteins within corona-nanoparticle complexes relative to their source fluid was determined *via* LC-MS analysis and is visually represented in Figure 3.

**Figure 3:**
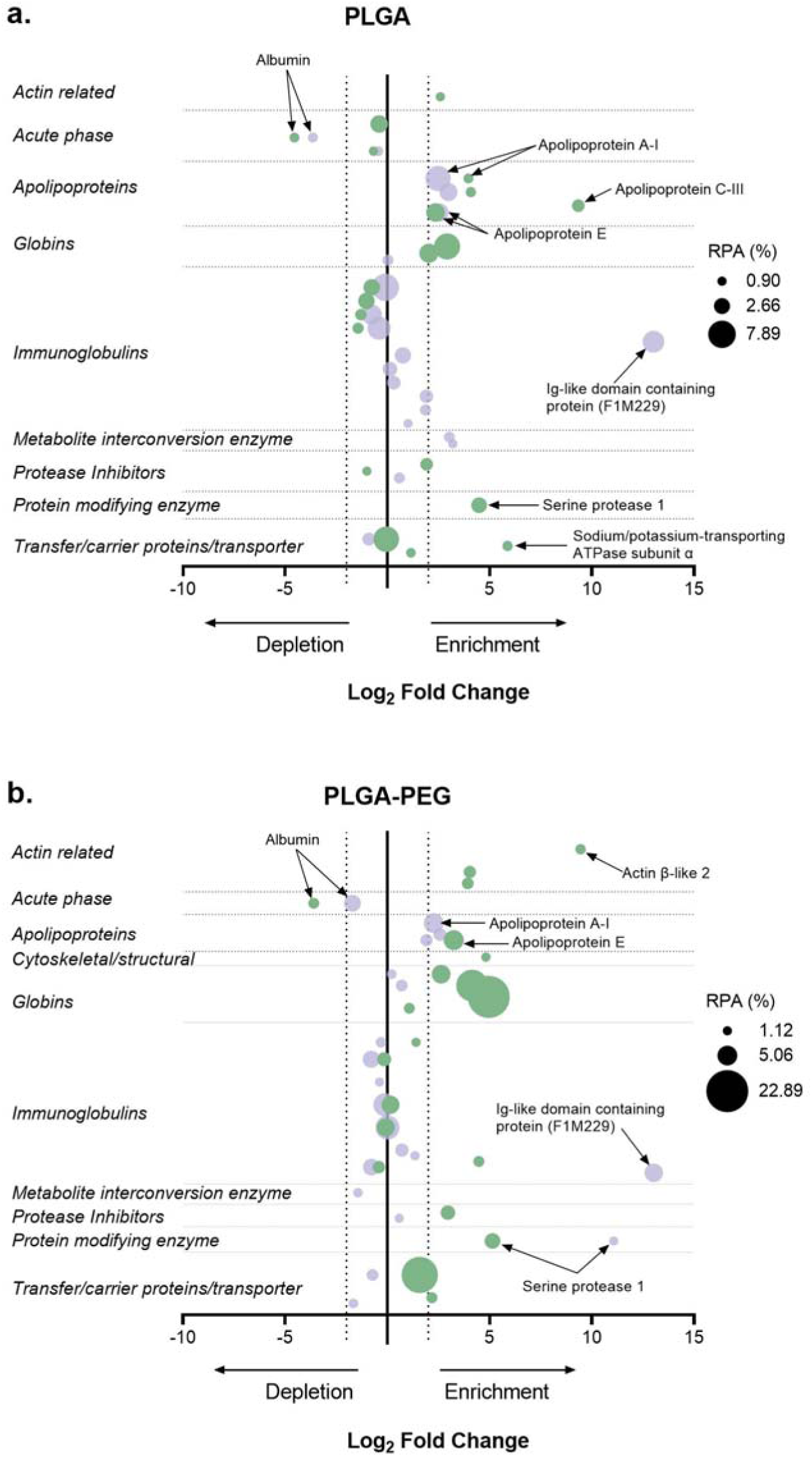
Compositional map illustrating the relative enrichment/depletion of serum (purple) or CSF (green) proteins in protein coronas formed on **(A)** PLGA or **(B)** PLGA-PEG nanoparticles. Circle size denotes the relative protein abundance within the top 200 proteins (RPA %) determined by LC-MS analysis. Proteins were categorized into functional classes using the PANTHER classification system (or UniProtKB). Log_2_ fold-change is in comparison/relation to the source biological fluid alone, where < 0 = depletion; > 0 = enrichment.

**Table 1:**
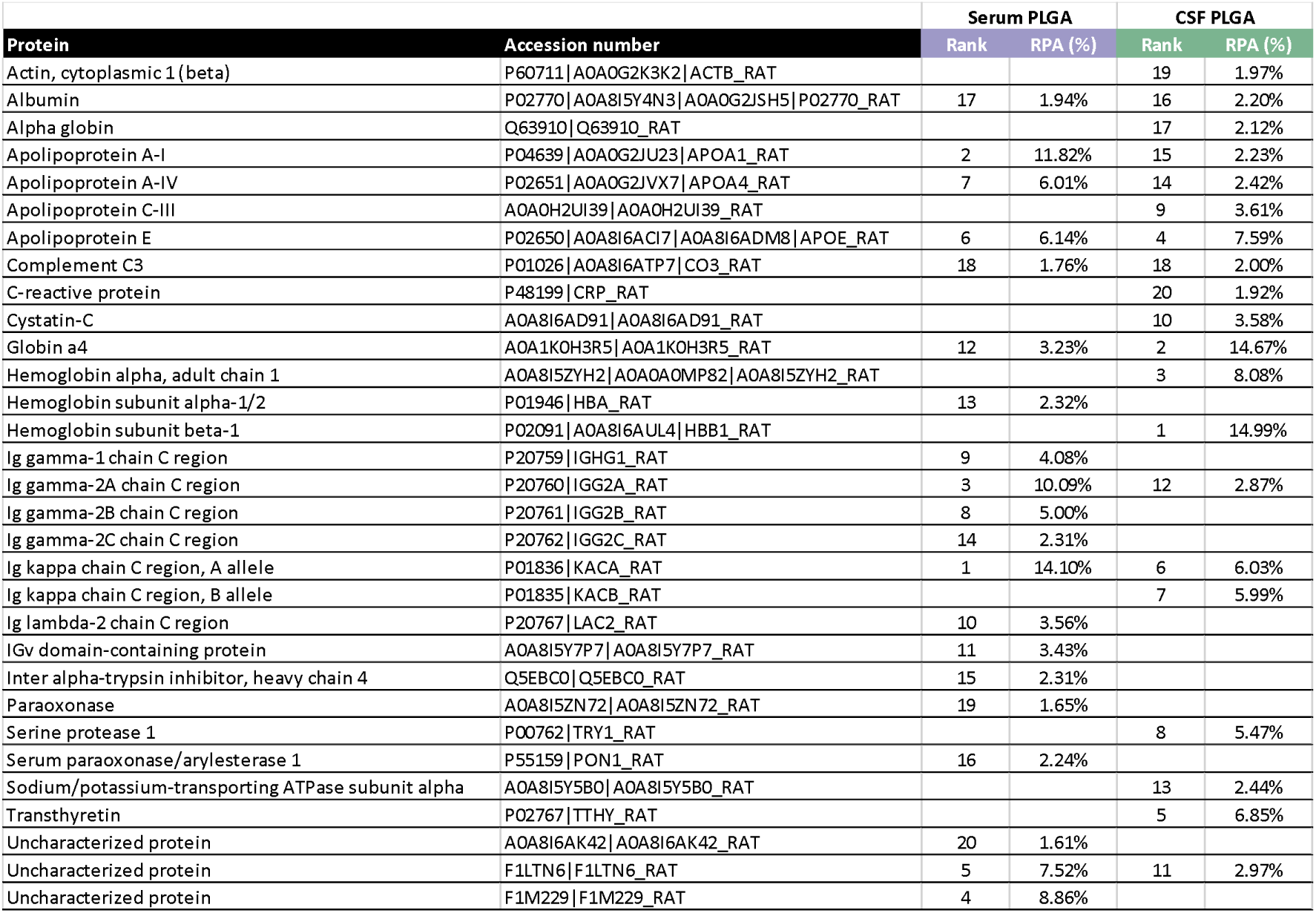
The Top-20 proteins adsorbed onto the surface of the PLGA nanoparticles serum- and CSF- and their relative protein abundance (RPA) (alphabetical order).

**Table 2:** The Top-20 serum (left) or CSF (right) proteins adsorbed onto the surface of the PLGA-PEG nanoparticles and their relative abundance (alphabetical order).

| Protein | Accession number | Serum PLGA-PEG |  | CSF PLGA-PEG |  |
| --- | --- | --- | --- | --- | --- |
|  |  | Rank | RPA (%) | Rank | RPA (%) |
| Actin, beta-like 2 | D3ZRN3\|D3ZRN3_RAT |  |  | 18 | 1.52% |
| Actin, cytoplasmic1 (beta) | P60711\|A0A0G2K3K2\|ACTB_RAT |  |  | 12 | 2.08% |
| Albumin | P02770\|A0A8I5Y4N3\|A0A0G2JSH5\|P02770_RAT | 7 | 6.68% | 16 | 1.64% |
| Alpha globin | Q63910\|Q63910_RAT |  |  | 14 | 1.68% |
| Apolipoprotein A-I | P04639\|A0A0G2JU23\|APOA1_RAT | 3 | 9.20% |  |  |
| Apolipoprotein A-IV | P02651\|A0A0G2JVX7\|APOA4_RAT | 9 | 4.05% |  |  |
| Apolipoprotein E | P02650\|A0A8I6ACI7\|A0A8I6ADM8\|APOE_RAT | 10 | 3.56% | 4 | 5.48% |
| Cystatin-C | A0A8I6AD91\|A0A8I6AD91_RAT |  |  | 9 | 2.86% |
| Globin a4 | A0A1K0H3R5\|A0A1K0H3R5_RAT | 12 | 3.30% | 2 | 17.77% |
| Hemoglobin alpha, adult chain 1 | A0A8I5ZYH2\|A0A0A0MP82\|A0A8I5ZYH2_RAT | 14 | 2.37% | 5 | 4.75% |
| Hemoglobin subunit beta-1 | P02091\|A0A8I6AUL4\|HBB1_RAT | 11 | 3.33% | 3 | 13.57% |
| Hemoglobin subunit beta-2 | P11517\|HBB2_RAT |  |  | 1 | 23.91% |
| Hemoglobin subunit zeta | G3V8R3\|G3V8R3_RAT |  |  | 17 | 1.62% |
| Hemopexin | P20059\|HEMO_RAT | 16 | 2.20% |  |  |
| IF rod domain-containing protein | A0A0G2JUG1\|A0A0G2JUG1_RAT |  |  | 19 | 1.23% |
| Ig gamma-1 chain C region | P20759\|IGHG1_RAT | 13 | 2.70% | 20 | 1.21% |
| Ig gamma-2A chain C region | P20760\|IGG2A_RAT | 5 | 6.96% | 10 | 2.72% |
| Ig gamma-2B chain C region | P20761\|IGG2B_RAT | 20 | 2.03% |  |  |
| Ig kappa chain C region, A allele | P01836\|KACA_RAT | 2 | 12.50% | 7 | 4.44% |
| Ig kappa chain C region, B allele | P01835\|KACB_RAT | 1 | 13.66% | 6 | 4.49% |
| Ig lambda-2 chain C region | P20767\|LAC2_RAT | 8 | 4.23% |  |  |
| IgV domain-containing protein | A0A8I5Y7P7\|A0A8I5Y7P7_RAT | 17 | 2.10% |  |  |
| Immunoglobulin heavy constant epsilon | F1LN61\|F1LN61_RAT |  |  | 15 | 1.66% |
| Inter alpha-trypsin inhibitor, heavy chain 4 | Q5EBC0\|Q5EBC0_RAT | 18 | 2.07% |  |  |
| Serine protease 1 | P00762\|TRY1_RAT | 19 | 2.06% | 8 | 3.35% |
| Serotransferrin | P12346\|TRFE_RAT | 15 | 2.28% |  |  |
| Similar to Actin, cytoplasmic 2 (Gamma-actin) | A0A8I5Y704\|A0A8I5Y704_RAT |  |  | 13 | 1.89% |
| Uncharacterized protein | F1LTN6\|F1LTN6_RAT | 6 | 6.69% | 11 | 2.13% |
| Uncharacterized protein | F1M229\|F1M229_RAT | 4 | 8.04% |  |  |

Among the Top-20 proteins, albumin emerged as the most abundant in both native serum (18%) and CSF (28%) (**Supplementary Table S3**). However, its adsorption to PLGA (**Figure 3a**) or PLGA-PEG particles (**Figure 3b**) was notably limited, with PLGA coronas showing a 10-fold or 13-fold depletion, and PLGA-PEG particles showing 3-fold or 17-fold depletion, compared to the source serum or CSF, respectively. This finding, at first, seems to contradict with prior observations of CSF-derived corona formation on PEGylated polystyrene particles (PEG-PNPs), where albumin accounts for ∼36-39% of the proteins in the total corona.^[15]^ In contrast, it is in agreement with research on serum- and CSF-derived corona formation on non-PEGylated particles and carbon nanotubes, where albumin is over 2.5 million-fold lower as compared to source fluid.^[14]^ Albumin is a notable high abundance-low affinity protein^[27]^ and readily binds to particles within a short time frame, such as the 10 minutes allowed for corona formation on PEG-PNPs in the previous study.^[15]^ This transient corona is then replaced by lower abundance, but higher affinity proteins overtime,^[31]^ as reported in the 1 hour corona formation on carbon nanotubes,^[14]^ and as we report here, where corona formation was allowed to occur over 6 hours.

The most abundant proteins in the serum PLGA corona (**Table 1**) were Ig Kappa chain C region, A allele (14%), apolipoprotein A-I (APOA1; 12%) and Ig gamma-2A chain C region (10%). Conversely, the most abundant proteins in the CSF PLGA corona were hemoglobin subunit beta-1 (15%), Globin a4 (15%), and hemoglobin subunit alpha (8%). This pattern was mirrored in the PLGA-PEG coronas (**Table 2**), where Ig Kappa chain C region, B allele (14%), Ig Kappa chain C region, A allele (13%) and APOA1 (9%) were the most abundant proteins in the serum PLGA-PEG corona, and hemoglobin subunit beta-2 (24%), Globin a4 (18%), and hemoglobin subunit beta-1 (14%) were the most abundant proteins in the CSF PLGA-PEG corona. The most depleted protein regardless of biological fluid or particle type was consistently albumin.

Evidently, the two serum coronas exhibited remarkably similar protein compositions, with immunoglobulins and apolipoproteins accounting for the most abundant functional classes (Figure 2e). This similarity extended to the Top-20 most abundant proteins, with the two serum coronas sharing 14 out of 20 proteins (**Table 1**, **Table 2**). Reduction in abundance of albumin and increased abundance of immunoglobins and apolipoproteins as compared to serum is expected, as immunoglobulins and apolipoproteins are reported to have higher affinity than albumin.^[27, 32]^

CSF corona compositions on both particles are similarly resemblant of each other sharing 13 of the Top 20 proteins, with globins accounting for the top 3 most abundant proteins on both. However, it must be noted that the number of detected proteins within the total coronas was wildly different; PLGA – >1000 proteins detected, PLGA-PEG – 28 proteins detected. As a result, the top 20 proteins reported in table 1 account for 37% of proteins within the total PLGA corona (47% within the top 200), while the top 20 proteins reported in table 2 account for 90% of proteins within the total PLGA-PEG corona. Regardless, this prevalence of globins in CSF-derived coronas aligns with previous findings reported for CSF coronas formed for only 10 mins *in vitro* and *in vivo* on polystyrene nanoparticles where globins were detected, albeit to a lesser degree.^[15]^ While it seems that both particles preferentially adsorb globins (17- and 40-fold greater, for PLGA and PLGA-PEG coronas, respectively, than the CSF control), it is likely mediated by different properties of the particles. Work using gold nanoparticles revealed a 4.5-fold higher binding affinity of hemoglobin to small nanoparticles (40 nm) as compared to larger ones (70 nm), ^[33]^ which may partially explain adsorption to the smaller PLGA-PEG particles (72 nm). Further, hemoglobin readily adsorbs to negatively charged nanoparticles,^[34–35]^ which may further increase its affinity for the negatively charged PLGA-PEG particles (-34 mV).

Despite similarities of adsorbed proteins between coronas formed in the same source fluid, there are distinct differences of adsorbed proteins on the same particle formed in the two different fluids. These results validate a recent assertion that the protein corona is transitional, and substantially changes in composition after transitioning from “blood” to “brain”.^[7]^

Interestingly, high abundance did not always equate to increased enrichment. Enrichment was defined here as a Log_2_ fold change (as compared to source fluid) greater than 2 (Log_2_ > 2). This method of evaluation of enrichment of proteins, rather than abundance, allows for a more robust determination of how the formed corona changes comparatively to the baseline (i.e. the native source fluid).^[36]^ In the serum PLGA corona, neither the 1^st^ nor 3^rd^ most abundant proteins (Ig kappa chain C region or Ig gamma 2-A chain C region, respectively) were enriched (**Figure 3a**). Ig-like domain containing protein (Uncharacterized protein; F1M229), the 4^th^ most abundant protein, was the single most enriched protein (Log_2_=13; Figure 3a). This is consistent with earlier reports, where immunoglobulins were depleted in serum coronas on polystyrene nanoparticles (PNPs) and carbon nanotubes.^[14]^ Additionally, apolipoproteins were the most enriched functional class of proteins, accounting for 3 out of 6 enriched proteins; namely, apolipoprotein A-IV (APOA4) (Log_2_=3; 7^th^ most abundant), apolipoprotein E (APOE) (Log_2_=2.5; 6^th^ most abundant), and apolipoprotein A-I (APOA1; Log_2_=2.5; 2^nd^ most abundant). While APOE and APOA4 have been shown to be enriched in serum coronas on polystyrene nanoparticles, APOA1 was depleted; conversely, on carbon nanotubes AOPE was enriched, while APOA1 and APOA4 were depleted.^[14]^ This highlights how nanoparticle characteristics can influence the specific protein enrichment in the protein corona, which may, in turn, have implications for distinct functional and safety profiles between different classes of nanoparticles.

In the CSF, PLGA corona apolipoproteins presented as the most enriched functional class, accounting for 4 out of 9 enriched proteins **(Figure 3a)**. Apolipoprotein C-III (APOC3) was the most strongly enriched protein (Log_2_=9.3), despite being only the 9th most abundant. Although the serum PLGA corona showed a remarkable enrichment of APOC3 (Log_2_=10), it only represented 0.6% of the Top-200 proteins (thus was outside of the Top 20 detected proteins in the serum PLGA corona). Specific apolipoproteins enriched in the CSF corona on PLGA particles were APOE (Log_2_=2.3; 4^th^ most abundant) and to a much greater degree, APOA1 (Log_2_=4; 15^th^ most abundant) and APOA4 (Log_2_=4; 14^th^ most abundant). This contrasts with earlier work, where apolipoproteins are consistently depleted, or not enriched, in CSF coronas on PNPs and carbon nanotubes.^[14]^ Other work shows depletion of APOE in CSF coronas on PEG-PNPs; however, enrichment/depletion was not specifically reported, nor was abundance of proteins outside the top 20 of CSF, so enrichment cannot be calculated for other apolipoproteins.^[15]^ This again reinforces how the choice of nanoparticle, and therefore its associated characteristics, will influence the composition of the adsorbed proteins corona.

Other greatly enriched Top-20 proteins in the CSF-PLGA corona were sodium/potassium-transporting ATPase subunit alpha (ATP1A; Log_2_=6; 13^th^ most abundant) and serine protease 1 (Log_2_=4.5; 8th most abundant). Greatly enriched

Top-20 proteins in Serum-PLGA coronas included paraoxonase (Log_2_=3.2;19^th^ most abundant) and serum paraoxonase/arylesterase (Log_2_=3; 16^th^ most abundant).

The serum PLGA-PEG corona showed a similar enrichment profile to its PLGA counterpart, with Ig-like domain containing protein the most enriched (Log_2_=13; 4^th^ most abundant; **Figure 3b**), and apolipoproteins accounting for 50% of enriched proteins; APOA1 (Log_2_=2.3; 3^rd^ most abundant) and APOA4 (Log_2_=2.6; 9^th^ most abundant). Conversely, only one apolipoprotein, APOE, was enriched on the CSF PLGA-PEG corona (Log_2_=3.2; 4^th^ most abundant). The most enriched functional classes in the CSF PLGA-PEG corona were globins and actin-related proteins, with 25% of enriched proteins falling into each of these classes.

Other greatly enriched proteins in the CSF PLGA-PEG corona were actin beta-like 2 (Log_2_=9.5; 18^th^ most abundant), and serine protease 1 (Log_2_=5.1; 8^th^ most abundant), while only serine protease 1 was additionally enriched in the serum PLGA-PEG corona (Log_2_=11; 19^th^ most abundant).

These distinct profiles suggest there are complex selective enrichment mechanisms driving protein adsorption.

### CSF coronas enhance the association of nanoparticles and brain cells

Protein coronas dictate the interactions between nanoparticles and cell membranes, and their subsequent mechanism and efficiency of cellular associations.^[37–38]^ We therefore set out to investigate how the different serum- and CSF-derived protein coronas influence the internalization of polymeric nanoparticles by a set of representative brain cell subtypes: brain endothelial cells; microglia; astrocytes; and differentiated neurons. In cellular uptake experiments (**Figure 4 and Supplementary Figure S2**), each cell type was incubated with FITC-loaded nanoparticles (0.5 mg/mL) both with and without coronas for a duration of 24 h, with the percentage of live cells containing fluorescent particles quantified by flow cytometry and visually confirmed *via* confocal fluorescence microscopy.

**Figure 4:**
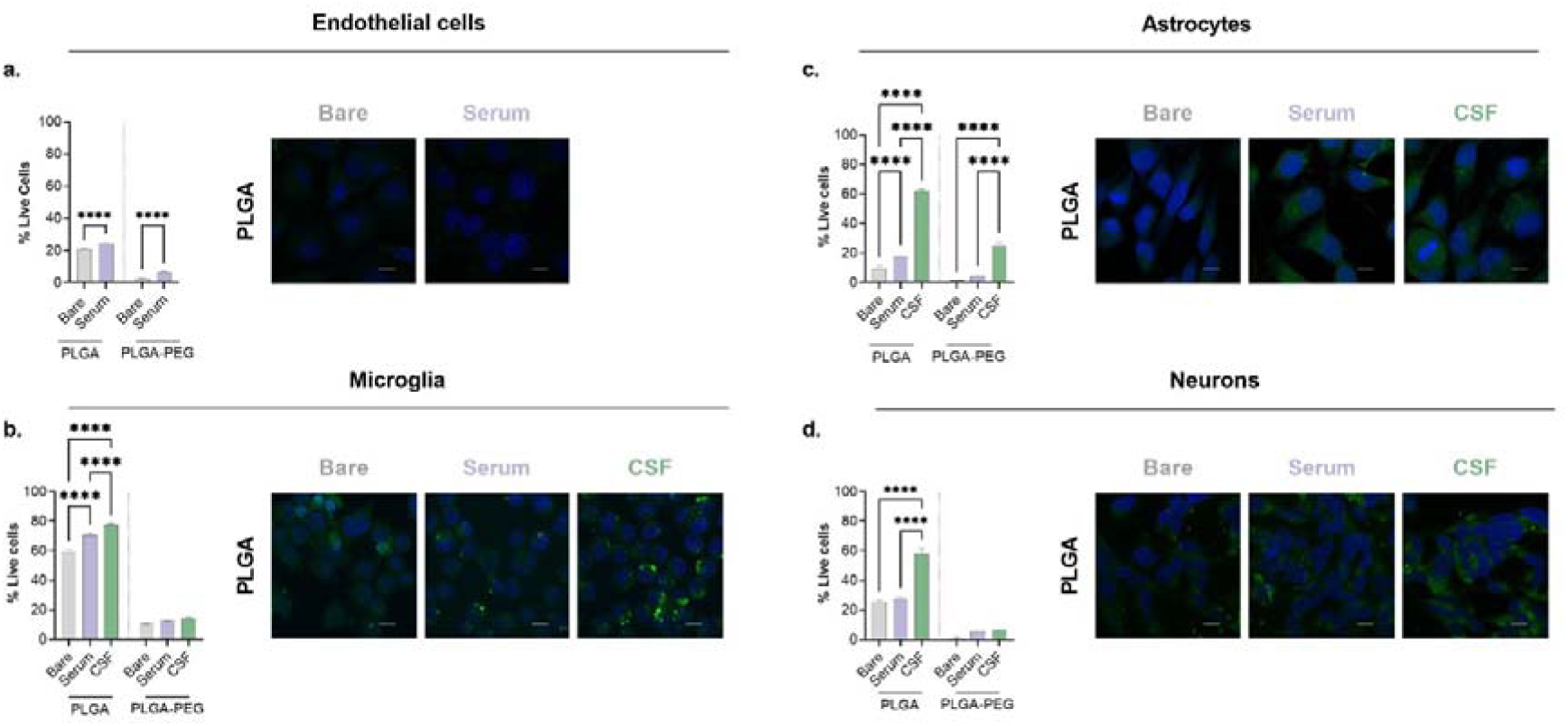
Cellular uptake of corona-nanoparticle complexes by different brain cell types. Fluorescent FITC-loaded PLGA or PLGA-PEG particles (0.5 mg/mL) were coated with serum or CSF coronas and incubated for 24 h with: **(a)** bEnd.3 brain endothelial cells; **(b)** BV2 microglia; **(c)**1321N1 astrocytes; and **(d)** SH-SY5Y-derived neurons. **(Left panel)** The internalization of fluorescent nanoparticles by live cells was quantified using flow cytometry. Error bars represent the mean ± standard deviation, two-way ANOVA, Šídák, n ≥ 3, from three independent experiments (**** p ≤ 0.0001). **(Right panel)** Merged confocal fluorescence microscopy images visualizing the cellular uptake of PLGA particles (green) with and without corona coatings; DAPI-stained cell nuclei (blue); Scale bars = 10 μm.

During their journey to the brain, systemically administered neuronanomedicines acquire serum coronas that will subsequently influence their capacity to penetrate the BBB. To reflect this delivery route, the effect serum-derived coronas had on the association of polymeric nanoparticles was initially assessed in bEnd.3 murine brain endothelial cells, which mimic various structural and functional properties of the BBB. In **Figure 4a**, 21% of bEnd.3 cells associated with bare PLGA particles, which increased slightly to 24% with serum coronas. On the other hand, poor cellular uptake was observed for PLGA-PEG both with and without serum coatings (< 7%).This is in line with previous literature, showing that brain endothelial cells demonstrate lower nanoparticle uptake efficiency than other endothelial cell types in the body, potentially reflecting the selective uptake nature of the BBB.^[39–40]^ This is due, in large part, to phenotypic differences between brain endothelial cells and endothelial cells of other vascular beds (e.g. lipid composition, presence of tight junctions),^[41]^ which affect BBB permeability and molecule transport.^[42]^

After crossing the BBB, neuronanomedicines enter the brain parenchyma, where the CSF facilitates biodistribution and subsequent engagement with glial cells (e.g., microglia, astrocytes) and neurons. Accordingly, corona-nanoparticle complexes were first incubated with microglia, brain-resident macrophages that play critical roles in innate immune responses and are key participants in neuroinflammation.^[43]^ Here, BV2 murine microglial cells readily phagocytosed bare PLGA nanoparticles, but were less efficient at internalizing PLGA-PEG particles (**Figure 4b**). Interestingly, the uptake efficiency of PLGA particles increased 21% and 33% with serum and CSF coatings, respectively. While corona-bearing PLGA-PEG particles did show a slight increase in uptake, their overall uptake remained low (< 10%). This underscores the purpose of PEGylation, which reduces protein corona formation on nanoparticles to lower immune cell recognition.^[6, 44]^ In previous reports, other polymeric nanoparticles injected directly into mice brains have shown a preferential internalization by microglia.^[45]^ This is not surprising, given the high phagocytic ability of microglia, with evidence that formation of the protein corona can further enhance this by activating phagocytic pathways (in a macrophage cell line).^[46]^ This is also in line with recent research that demonstrated that formation of a protein corona on the surface of fluorinated PLGA-based nanoparticles in cell culture medium (DMEM with 10% fetal bovine serum (FBS), 100 mg ml^−1^ of streptomycin, 100 U ml^−1^ of penicillin and 2 mM of l-glutamine) plays a key role in enhancing microglial internalization of these particles.^[47]^

Astrocytes, the most abundant and versatile glial cells in the CNS, provide structural and metabolic support to neurons.^[48]^ Model astrocytes, 1321N1 human astrocytoma cells, exhibited minimal uptake bare PLGA nanoparticles (< 10%), but this increased by ∼2-fold with serum coatings and almost 7-fold with CSF coatings (**Figure 4c**). Moreover, the internalization of PLGA-PEG particles without and with serum coronas was negligible (< 5%), however, CSF-derived coronas led to a remarkable 22-fold enhancement in astrocyte uptake. Overall, it was perhaps surprising that astrocytes showed such minimal uptake of the bare nanoparticles, given that previous literature has shown that the 1231N1 human astrocytoma cell line does endocytose other polymeric nanoparticles types through clathrin-mediated mechanisms.^[49]^ It must be noted, however, that, for the current work, astrocytes were not pre-stimulated with an inflammatory challenge (e.g. with lipopolysaccharide (LPS)). Stimulation of astrocytes transforms them to a reactive phenotype, which may affect their phagocytic capacity (for review, see Konishi et al. 2022).^[50]^ Thus, it is possible that different uptake results would have been seen under pro-inflammatory conditions, which also might be more representative of what is seen in the brain in many neurological disorders.^[51]^ This may also, at least in part, help to explain why increased uptake was seen with protein corona formation, at least for PLGA nanoparticles. As discussed above, there is evidence from a macrophage cell line that protein corona formation leads to activation of phagocytic pathways and subsequent enhanced nanoparticle uptake.^[46]^ While, to our knowledge, the effect of protein corona formation on uptake in an astrocyte cell line has not yet been evaluated, it is reasonable to hypothesize that it may have similar effects. Alternatively, it has been suggested that astrocytes will compensatively phagocytose dead cells in the absence of microglia.^[52]^ Therefore, it is possible that the astrocytes recognize some component within the CSF corona, which is not present in the serum corona, that can activate this phagocytic astrocyte mechanism.^[50]^

Neurons are the conducting cells of the CNS, and thus, the key players in functional brain activity. In this study, SH-SY5Y human neuroblastoma cells were differentiated into neurons (**Supplementary Figure S1**) and treated with corona-nanoparticle complexes (**Figure 4d**). PLGA particles without or with serum coatings exhibited similar uptake efficiencies of ∼25%, while CSF-derived coronas elevated uptake by over 2-fold. Under all conditions tested, neurons displayed a low capacity to take up PLGA-PEG particles (< 7%).

Collectively, these results reveal the ability of serum- and CSF-derived protein coronas to augment nanoparticle uptake across a range of brain cell subtypes. CSF coronas promoted internalization significantly more than their serum counterparts under all conditions tested, with substantial enhancements observed for CSF-coated PLGA particles interacted with astrocytes and neurons in particular. Similarly, despite PEGylated particles consistently displaying poor cellular internalization, CSF coronas greatly enhanced their uptake by astrocytes. Given that serum PLGA coronas showed enhanced uptake in microglia and astrocytes as compared to bare particles, but had no effect on the uptake of neurons, it may suggest that that the serum corona enhances phagocytic pathways in glial cells. Conversely, the CSF corona showed enhanced uptake in glial cells as well as neurons suggesting internalization by phagocytic and non-phagocytic pathways.

### Key proteins in CSF coronas correlate with brain cell uptake

To identify key proteins affecting the internalization of polymeric nanoparticles by brain cells, a Pearson’s correlation coefficient (*r*) analysis was conducted to elucidate relationships between protein corona composition and cellular uptake efficiency (**Table 3**).^[39]^

**Table 3:** Correlation analysis between nanoparticle uptake in brain-related cells and the relative abundance of corona proteins. Positive correlation coefficients (*r*) ≥ 0.6 are shaded green and negative correlation coefficients (*r*) ≤ 0.6 are shaded red. Brain endothelial cells were exposed to serum coated nanoparticles only, and therefore had too few pairs for correlation analysis.

| Protein | Microglia | Astrocytes | Neurons |
| --- | --- | --- | --- |
| Actin, beta-like 2 | -0.38 | -0.19 | -0.55 |
| Actin, cytoplasmic 1 (beta) | -0.20 | -0.03 | -0.39 |
| Albumin | -0.50 | -0.90 | -0.66 |
| Alpha globin | 0.11 | 0.37 | -0.03 |
| Apolipoprotein A-I | -0.44 | -0.51 | -0.24 |
| Apolipoprotein A-IV | -0.28 | -0.32 | -0.04 |
| Apolipoprotein C-III | 0.75 | 0.30 | 0.70 |
| Apolipoprotein E | -0.23 | 0.20 | -0.27 |
| Complement C3 | 0.35 | 0.19 | 0.54 |
| C-reactive protein | 0.89 | 0.56 | 0.87 |
| Cystatin-C | 0.12 | 0.35 | -0.04 |
| Globin a4 | -0.13 | 0.13 | -0.28 |
| Hemoglobin alpha, adult chain 1 | 0.23 | 0.27 | -0.01 |
| Hemoglobin subunit alpha-1/2 | -0.29 | 0.08 | 0.04 |
| Hemoglobin subunit beta-1 | 0.02 | 0.19 | -0.18 |
| Hemoglobin subunit beta-2 | -0.36 | -0.07 | -0.49 |
| Hemoglobin subunit zeta | -0.36 | -0.07 | -0.49 |
| Hemopexin | -0.43 | -0.73 | -0.37 |
| IF rod domain-containing protein | -0.36 | -0.07 | -0.49 |
| Ig gamma-1 chain C region | -0.83 | -0.61 | -0.62 |
| Ig gamma-2A chain C region | -0.78 | -0.61 | -0.56 |
| Ig gamma-2B chain C region | -0.33 | -0.19 | -0.04 |
| Ig gamma-2C chain C region | -0.32 | -0.31 | -0.07 |
| Ig kappa chain C region, A allele | -0.81 | -0.78 | -0.66 |
| Ig kappa chain C region, B allele | -0.31 | -0.73 | -0.57 |
| Ig lambda-2 chain C region | -0.46 | -0.68 | -0.35 |
| IGv domain-containing protein | -0.46 | -0.43 | -0.22 |
| Immunoglobulin heavy constant epsilon | -0.64 | -0.32 | -0.71 |
| Inter alpha-trypsin inhibitor, heavy chain 4 | -0.25 | -0.43 | -0.08 |
| Paraoxonase | -0.13 | -0.15 | 0.12 |
| Serine protease 1 | 0.13 | 0.24 | -0.09 |
| Serotransferrin | -0.22 | -0.59 | -0.17 |
| Serum paraoxonase/arylesterase 1 | -0.24 | -0.29 | -0.01 |
| Similar to Actin, cytoplasmic 2 (Gamma-actin) | -0.27 | -0.11 | -0.47 |
| Sodium/potassium-transporting ATPase subunit alpha | 1.00 | 0.81 | 0.93 |
| Transthyretin | 0.93 | 0.61 | 0.90 |
| Uncharacterized protein (F1LTN6) | -0.77 | -0.76 | -0.61 |
| Uncharacterized protein (F1M229) | -0.55 | -0.64 | -0.38 |
| Uncharacterized protein (A0A8I6AK42) | -0.43 | -0.55 | -0.25 |

A positive correlation (*r* ≥ 0.6), indicating that that particular protein is associated with cellular uptake, was observed in all three brain cell types for ATP1A1 and Transthyretin (TTY). Interestingly, APOC3 and C-reactive protein (CRP) showed a positive correlation in microglia and neurons, but no correlation in astrocytes. Overall, these positively correlated proteins accounted for 15% of the Top-20 most abundant proteins identified in CSF coronas on PLGA nanoparticles (**Table 1**). In contrast, none of the positively correlated proteins featured in the Top-20 of any other corona-nanoparticle complex (**Table 2** and **3**), therefore, these four CSF specific proteins may be involved in a mechanism resulting in consistent enhanced uptake of these CSF-coated PLGA particles.

APOC3 is primarily known for its role in lipid metabolism, rather than direct involvement in enhancing brain cell uptake of nanoparticles. The enhancement of brain cell uptake is more commonly associated with other apolipoproteins, like APOE, the most highly expressed apolipoprotein in the brain,^[53]^ which has a well-established role in mediating the transport of nanoparticles across the BBB.^[54–56]^ Moreover, internalization into glial cells is likely mediated by APOE *via* low density lipoprotein (LDL)-receptor mediated transcytosis,^[3]^ consistent with the opsonizing effects of apolipoproteins.^[57]^ Apolipoprotein mediated transport appears to be apolipoprotein-, particle- and/or cell type-dependent, since it has been reported that polystyrene nanoparticles precoated with APOA4 or APOC3 exhibited significantly decreased cellular uptake, whereas nanoparticles precoated with apolipoprotein H displayed increased uptake in human mesenchymal stem cells.^[58]^ Conversely, in a different study, of all apolipoproteins tested, only APOC3 facilitated internalization of lipid-based nanoparticles into mast cells, potentially through APOE receptor 2 binding.^[59]^ Curiously, since astrocytes and microglia are derived from the same glial precursors, it could be presumed that internalization occurs through a similar mechanism, despite apolipoproteins not showing correlation with uptake in astrocytes here. However, the field of nanomedicine is rapidly evolving, and novel roles for various apolipoproteins, including APOC3, in promoting cellular uptake might emerge with further research.

TTY is a transport protein, categorized here as a negative acute phase protein, as its levels decrease in response to inflammation.^[60]^ While it primarily transports retinol and thyroxine, it has also demonstrated BBB transcytosis capabilities. TTY conjugated to quantum dot nanoparticles was shown to facilitate transcytosis across endothelial cells *via* clathrin mediated transcytosis *in vitro,* and enhanced brain distribution of particles following intravenous administration *in vivo.*^[61]^ Further, TTY has been shown to transport amyloid beta across BBB endothelial cells (from brain to blood, mono-directionally) *via* the LDL receptor-related protein 1 (LRP1).^[62]^ However, here, despite not showing enrichment in the CSF PLGA corona (Log_2_=- 0.4), TTY accounted for 7% of top 20 corona composition suggesting, perhaps, a moderate effect on internalization.

ATP1A1 functions as an ion transporter to pump sodium out of and potassium in to cells at the expense of ATP. ATP1A1 is ubiquitously expressed across neuronal cell membranes,^[63–64]^ as well as in BBB endothelial cells.^[65]^ While it is unlikely that ATP1A1 itself is involved in internalization of nanoparticles, its activation has been shown to be crucial for macropinocytic internalization of respiratory syncytial virus in lung epithelial cells.^[66]^ ATP1A1 activation resulted in downstream epidermal growth factor receptor (EGFR) transactivation, leading to the formation of macropinosomes. This suggests that activation of EGFR on cells by ATP1A1 in the CSF corona could contribute to clathrin-mediated or -independent endocytosis.^[67]^ In line with this, EGFR has previously been shown to bind to multiple pathological protein aggregates implicated in neurodegenerative disease, including amyloid beta,^[68]^ huntingtin and alpha-synuclein,^[69]^ mediating their uptake into cells.

Lastly, CRP is a positive acute phase protein and stimulates indirect phagocytosis through complement activation, and direct receptor-mediated phagocytosis by enhancing IgG binding, or directly binding to the Fc gamma receptors (FcγR) on immune cells, thereby acting as an opsonin.^[70–71]^ Microglia express all 3 types of FcγRs (CD16, CD32 and CD64),^[72]^ while neurons and astrocytes primarily express the high affinity FcγR CD64. ^[73–74]^ Interestingly, immunoglobulins, which bind FcγRs, were not positively correlated, and many were, in fact, negatively correlated with uptake across all 3 cell lines. Indeed, the serum-coated PLGA particles showed less internalization into microglia than CSF-coated PLGA particles, despite immunoglobulins making up 36% of the composition compared to 10%. Further, neither immunoglobulins nor CRP were enriched in the CSF PLGA corona, suggesting that CRP-mediated phagocytosis was minimal.

Taken together, these proteins individually, or in conjunction with each other, may contribute to enhanced internalization of CSF corona coated PLGA nanoparticles, although the exact mechanisms of this remain to be explored.

This suggests that pre-adsorption of proteins identified as enriched in the CSF PLGA corona, or other CSF-specific enriched proteins, to the surface of nanoparticles could be used to manipulate cell targeting through enhancing specific uptake mechanisms.^[75–76]^ Pertinently, pre-adsorption of beta-amyloid proteins to the surface of liposomes has been shown to increase the adsorption of apolipoproteins, leading to their internalization into brain cells *via* LDL-receptor mediated uptake.^[56]^ Further, it is established that different cells internalize the same particle through different mechanisms.^[49]^ Therefore, internalization studies of pre-coated nanoparticles in tandem with transport inhibitors could elucidate the effect of these proteins and their method of internalization.

Conversely to the positive associations discussed above, a negative correlation (*r* ≤ −0.6) associates a protein present in a corona with reduced cellular uptake. This was seen in all three brain cell types for Ig gamma-1 chain C region, Ig kappa chain C region (A allele) and Ig-like domain-containing protein F1LTN6. Moreover, albumin exhibited a negative correlation with particle internalization in astrocytes and neurons, but no correlation in microglia; while Ig gamma-2A chain C region correlated with decreased uptake in microglia and astrocytes, and no correlation in neurons. Together, these negatively correlated proteins constituted over 37% of the Top-20 proteins identified in serum coronas and less than 15% in CSF counterparts (**Table 1** and **2**).

As reported above, astrocytes demonstrated the lowest uptake, except for CSF PLGA particles. Unsurprisingly, a number of additional proteins showed a negative correlation in astrocytes only: Ig kappa chain C region (B allele), Ig lambda-2 chain C region, Ig-like domain-containing protein F1M229, and Hemopexin.

Notably, of the 10 proteins identified to be negatively correlated with uptake in any cell line, 8 of them were classified as immunoglobulins, which corresponded to 36% of the top 20 protein composition of both the serum PLGA and PLGA-PEG coronas, 16% of the CSF PLGA-PEG corona, but only 10% of the CSF PLGA corona. Immunoglobulins are opsonins and play a role in the adaptive immune response, as discussed above, by binding to the Fcγ receptors on phagocytic cells. Therefore, considering their abundance, it is unexpected that immunoglobulins are not correlated with uptake, particularly in the microglia. However, a study investigating the protein corona on PVA-PLGA and PVA-PLGA-PEG nanoparticles suggested that, while there may be an abundance of opsonins, dysopsonins may be covering their active sites, preventing or limiting phagocytosis.^[77]^ Therefore, further characterization into protein conformation/spatial interaction of dysopsonins and opsonins should be investigated.

While this correlation analysis gives interesting insights into potential protein-cell relationships, taken alone, it is insufficient to conclude causative mechanisms. In recent similar work, Alyandi *et al.* investigated the corona composition on silica nanoparticles and their uptake into organ-specific endothelial cells.^[39]^ Initial correlation analysis identified proteins associated with uptake (e.g. serotransferrin), however, when nanoparticles were coated with single-protein coronas, uptake was unchanged as compared to uncoated nanoparticles. When probed further, it was determined that uptake was likely by multiple mechanisms. Therefore, additional experiments to probe transport mechanisms such as single protein coronas and receptor inhibition, as well as further robust linear regression analysis, should be undertaken.

### CSF coronas reduce nanoparticle cytotoxicity and induce pro-inflammatory cytokine responses in brain cells

The protein corona influences cellular and immune responses towards nanoparticles (e.g., neuroinflammation), thereby affecting the immunotoxicity and safety of potential neuronanomedicines. In this study, flow cytometric analysis was used to measure the viability of brain cell subtypes after exposure to polymeric nanoparticles (with and without coronas) at different concentrations (0, 0.5, or 1 mg/mL) (**Supplementary Figure 3**). With the exception of neurons, all tested cells showed no change in viability, which is expected, considering both PLGA and PLGA-PEG are FDA-approved as safe and non-toxic polymers.^[18]^ Bare PLGA nanoparticles were found to significantly reduce the viability of neurons at higher concentrations (**Supplementary Figure 3)**. Nevertheless, this apparent cytotoxicity was alleviated when particles were coated with serum or CSF coronas.

To compare the effects serum- and CSF-derived protein coronas have on the immune responses to polymeric nanoparticles, brain cell subtypes were treated with corona-nanoparticle complexes (0.5 mg/mL). After 24 h, the secretion of proinflammatory cytokines, IL-6 and TNFα, were assayed *via* cytokine bead array (**Figure 5**). IL-6 regulates immune responses and metabolic processes, whereas TNFα is a potent proinflammatory cytokine that contributes to the initiation and amplification of inflammation.^[43, 78]^

**Figure 5:**
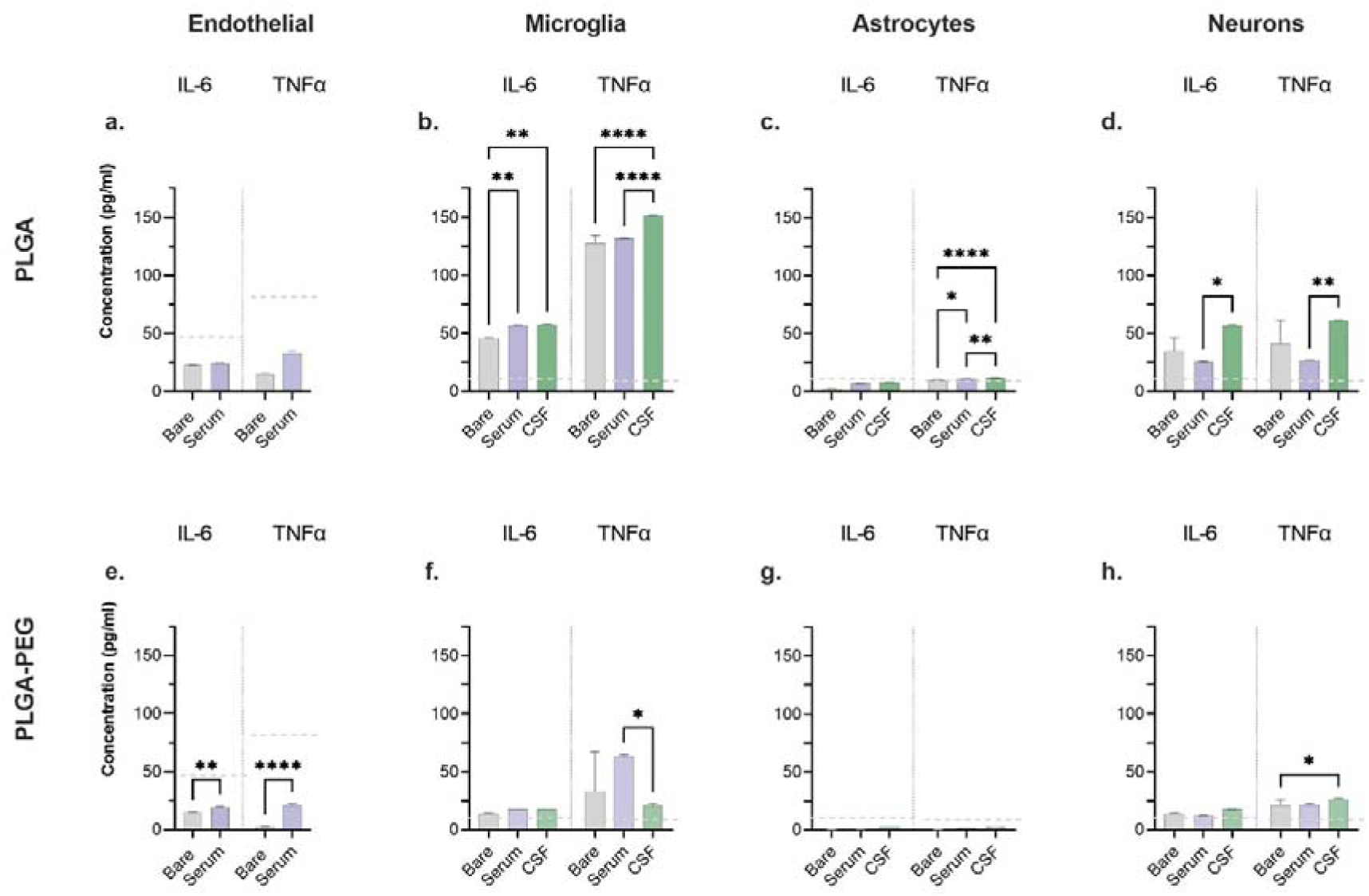
Inflammatory responses of brain cell subtypes to corona-nanoparticle complexes. **(Upper panel)** PLGA or **(Lower panel)** PLGA-PEG particles were coated with hard coronas derived from serum or CSF, and then incubated with: **(a)** bEnd.3 endothelial cells **(b)** BV2 microglia; **(c)** 1321N1 astrocytes; or **(d)** SH-SY5Y-derived neurons. 24 h after treatment, the concentration (pg/mL) of pro-inflammatory cytokines IL-6 and TNF-α released by cells was measured *via* cytokine bead array. Error bars represent the mean ± SD (*p ≤ 0.05, ** p ≤ 0.01, **** p ≤ 0.0001), two-way ANOVA, Šídák’s post-hoc analysis, n≥3, from three independent experiments. Dashed lines represent thresholds above normal levels for IL-6 or TNF. Grey = Polymeric nanoparticle; Purple = Serum; Green = CSF. Dotted lines represent normal physiological concentrations.

When exposed to polymeric nanoparticles without or with serum coronas (**Figure 5a, e**), bEnd.3 cells released cytokines at concentrations well below those normally found in the bloodstream (i.e., IL-6 ≤ 43 pg/mL; TNFα ≤ 75 pg/mL),^[79–80]^ further demonstrating the resilience of these cells towards nanoparticles.

In the brain, IL-6 and TNFα levels are ≤ 10 pg/mL and ≤ 6 pg/mL, respectively.^[81–83]^ Of the brain cell subtypes evaluated, microglia displayed the highest proinflammatory responses (**Figure 5b**). When treated with PLGA nanoparticles, BV2 microglial cells secreted 46 pg/mL of IL-6, which increased to ∼56 pg/mL with serum- and CSF-derived coronas. Moreover, both bare and serum-coated PLGA nanoparticles triggered the release of ∼130 pg/mL of TNFα, which elevated to ∼150 pg/mL with CSF coronas. Microglia exhibited relatively lower proinflammatory responses to PLGA-PEG nanoparticles, except for the serum-coated particles which induced the secretion of 63 pg/mL TNFα (**Figure 5f**). Together, these immune responses affirm the ability of microglia to phagocytose nanoparticles, and may suggest polarization to an M1 phenotype, due to the high levels of pro-inflammatory cytokines released.^[43]^

In contrast, astrocytes (**Figure 5c, g**) showed no significant proinflammatory responses to treatments, except for an elevation in TNFα levels after exposure to PLGA particles (both with and without coronas). These observations were somewhat surprising, given that astrocytes, much like microglia, contribute to innate immunity by producing cytokines.^[84]^ Low pro-inflammatory cytokine secretions were reasonably expected given the low level of uptake of the bare and serum-coated PLGA nanoparticles. However, the unexpectedly high internalization of CSF particles, and concomitant low secretion of pro-inflammatory cytokines, suggests that the CSF-corona didn’t stimulate activation of astrocytes, resulting in the production of pro-inflammatory cytokines. This assertation is supported by previous research where human astrocytes were incubated with various pro-inflammatory stimuli, however, their secretion of IL-6 was unchanged compared to unstimulated cells.^[85]^

Neurons showed varying proinflammatory responses under the conditions tested (**Figure 5d, h**). When coated with CSF coronas, PLGA nanoparticles induced significantly higher cytokine release than both the bare and serum-coated particles, a somewhat unsurprising outcome given that CSF-coated particles showed significantly higher uptake. This trend was similarly observed for PLGA-PEG particles, albeit with substantially lower cytokine levels overall.

These findings indicate that CSF-coated nanoparticles induce higher cell immune responses than bare or serum-coated nanoparticles, with comparatively lower cytokine release observed with PEGylated particles (with and without coronas). This is again unsurprising, as PEG coating is routinely used as an anti-fouling agent for the purpose of reducing immune interactions.^[22]^ Compared to corona-nanoparticle complexes, the cellular immune responses to bare nanoparticles tended to exhibit the highest variability. This variability was particularly evident when bare PLGA particles were interacted with neurons, which could partly explain the variations in neuron viability mentioned above (**Supplementary Figure S3**). Overall, cellular immune responses correlated to some extent with the respective cellular uptake of each corona-nanoparticle complexes shown earlier (**Figure 4**).

## 3. Conclusion

The formation of a protein corona is an inescapable consequence of nanomedicine delivery. Therefore, understanding interactions at the bio-nano interface, and the consequence of these interactions on nanoparticle-corona complexes and target cells, is essential. Insufficient understanding of these interactions likely contributes to the lack of clinical translation of neuronanomedicines. Our findings highlight the importance of recognizing both the similarities and differences in protein coronas formed in different biological and physiologically relevant fluids when evaluating neuro-nanomedicines. We have shown that distinct protein coronas are formed on particles in different biological fluids, and shown preferential internalization of a CSF-derived protein corona into brain-related cells. Moreover, we report, for the first time, the effect of the CSF protein corona on cellular cytokine release, with important implications for neuroinflammation and cytotoxicity of neuronanomedicines.

We have underscored the importance of appropriate *in vitro* model selection in one capacity only, ie. physiologically relevant biofluid. However, there are several other parameters which need to be considered in *in vitro* modelling for NDDS development. While our work and others^[15]^ have used traditional 2D cell culture models, previous studies have shown substantial protein corona changes after passing through a transwell BBB model.^[7]^ Recent findings by Lin *et al.* have also demonstrated a changed local environment in an injury model using endothelial cells transiently affected the composition of the protein corona.^[86]^ Further, it is also well established that corona formation is disease and sample specific.^[8, 87]^ Therefore investigation into the corona formed in diseased source fluids will inform the development of neuronanomedicines. Ultimately, the model chosen, reflecting the route of administration and the physiological environment, needs to be carefully considered in future *in vitro* CSF corona studies.

Recent comprehensive reviews of the literature have highlighted sources of variability and error in protein corona studies, which will affect their interpretation and future clinical translation.^[12, 88]^ Therefore, the standardization of protein corona methodologies and analysis is imperative to produce complete, robust and reproducible protein corona data, allowing for better understanding of bio-nano interactions, and leading to *in vivo* translation.

In addition, by drawing inspiration from small-molecule drug development, standardization of corona studies will allow for the generation of large and robust datasets. These datasets of nanoparticle-BBB interactions could be curated to create machine learning libraries and algorithms.^[89–90]^ This was recently applied to identify which proteins adsorb to carbon nanotubes. The random forest classifier model of machine learning identified adsorbed proteins based off protein sequence with 78% accuracy, 70% precision and 65% recall.^[91]^ Further, modelling has been used to predict the cytotoxicity of nanoparticles,^[92–95]^ predict the formation of a protein corona on a range of nanoparticles,^[96–97]^ and, importantly, predict the effect of the protein corona on cellular interactions^[97–99]^ and biological fate.^[100]^ In conjunction with *in silico* modelling techniques, these tools could optimize existing NDDSs or even predict entirely new designs capable of effectively crossing the BBB.

Armed with a better understanding, there is the potential to alter the physicochemical properties of nanoparticles (e.g., surface chemistry) to manipulate PC formation. This would enable more control over the biological fate of an NDDS (e.g., enhanced BBB crossing, pharmacokinetics, biodistribution, biocompatibility, and clearance), which could greatly improve a neuronanomedicine’s overall therapeutic activity.

## 4. Experimental Section

### Materials

PLGA-PEG-NH_2_ copolymer (Mw 12,000-2,000Da) was purchased from Akina Inc (USA). Acid terminated PLGA copolymer (Resomer RG 503’ lactide:glycolide 50:50; Mw 24,000-38,000Da); and poly(vinyl alcohol) (PVA; Mowiol 8-88; Mw ∼67,000Da) were both purchased from Sigma-Aldrich. Normal rat serum and pooled CSF collected from Sprague-Dawley rats were obtained from Invitrogen and BioIVT (USA), respectively. All other chemicals and reagents used in this study were purchased from Sigma-Aldrich or ThermoFisher Scientific, unless stated otherwise.

### Nanoparticle preparation and biophysical characterization

*Polymeric nanoparticle synthesis:* PLGA and PLGA-PEG nanoparticles were synthesized using an emulsion-evaporation method with polyvinyl alcohol (PVA) as a stabilizer, following a protocol adapted from Locatelli *et al..* ^[23]^ Briefly, 30 mg of either PLGA or PLGA-PEG copolymer was dissolved in dichloromethane (DCM) and added dropwise to a 5% PVA solution (12.5 mL), followed by vortexing. The mixture then underwent sonication on ice for a total duration of 1.5 min (3 x 30 s cycles) at 100% amplitude using a microtip-probe sonicator. The resulting NP-containing solution was placed in a shaker overnight to facilitate organic phase evaporation. Following this, nanoparticles were collected by centrifugation at 17,500 rcf for 15 min and redispersed in water. This washing process was repeated twice to remove any residual solvent/stabilizer from the particles. Finally, the purified particles were redispersed in 1 mL of ultrapure water and stored at 4°C until further use.

#### Fluorescent Dye Encapsulation in Nanoparticles

The green fluorescent dye, Fluorescein isothiocyanate isomer 1 (FITC), was loaded into PLGA and PLGA-PEG nanoparticles using a double emulsion-solvent evaporation method.^[23]^ 30 mg of the respective copolymer and 1 mg of FITC were co-dissolved in DCM. This mixture was added dropwise into 12.5 mL of 5% PVA and then vortexed before being subjected to sonication for 2 min total (4x 50 s cycles) at 100% amplitude on ice. The resulting solution, containing FITC-loaded nanoparticles, was stirred overnight in a beaker to evaporate the organic phase. Nanoparticles were collected *via* centrifugation, followed by resuspension in water. Any surplus dye was subsequently removed by overnight dialysis against PBS at 4°C, utilizing Slide-A-Lyzer Dialysis Cassettes with a 10 kDa molecular cut-off. The purified FITC-loaded nanoparticles were collected and stored at 4°C in the dark for use in downstream experiments.

#### Dye entrapment efficiency

To quantify the amount of FITC dye encapsulated by the polymeric nanoparticles, 30Cmg of FITC-loaded particles were suspended in DMSO. The sample was diluted in PBS and FITC concentrations were calculated using the fluorescence value obtained at 525 nm and a standard curve of FITC in methanol (0–1.6Cmg/mL). The percentage entrapment efficiency of the particles was then calculated as:

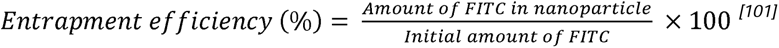

#### Dynamic light scattering (DLS)

The zeta-potential, PDI, and hydrodynamic diameter of nanoparticles (with and without corona coatings) were measured on a Zetasizer Nano ZS (Malvern Instruments, UK). Three measurements were performed at 25°C in cuvettes or capillary cells containing 1 mL of nanoparticles diluted in ultrapure water to a final concentration of 0.15 mg/mL. Data analysis was carried out using the Zetasizer Nano software.

#### Transmission electron microscopy (TEM)

To visualize nanoparticle size, morphology, and protein corona formation, TEM imaging was conducted with a JEOL JEM-F200 microscope operating at 120 kV. 10 μL of sample (0.1 mg/mL) were adsorbed overnight onto Pioloform-coated 200 mesh copper grids (ProSciTech). Prior to imaging, samples were negatively stained with 2% uranyl acetate replacement (UAR-EMS, Electron Microscopy Sciences) for 1 h, rinsed with ultrapure water, and then allowed to dry for at least 3 h.

### Protein corona formation and analysis

The formation of hard coronas on the nanoparticle surfaces was carried out according to a method modified from Partikel and co-workers.^[102–103]^ Here, suspensions of PLGA or PLGA-PEG particles with total surface areas (A = 4πr^2^) equivalent to 0.08 m^2^ were mixed with 75 μL of rat serum or CSF and incubated at 37°C for different time periods (15 and 30 min; 1, 3, 6, and 12 h). Following this, excess or weakly bound serum or CSF proteins were removed from the particles via three wash cycles of centrifugation (15 min, 21,000 rcf) and redispersal in 500 μL ultrapure water. Upon completion, purified corona-nanoparticle complexes were suspended in ultrapure water and stored at 4°C until further use.

#### Protein quantification

To quantify total protein adsorbed onto nanoparticles, corona-nanoparticle complexes were first desorbed from nanoparticles and denatured by incubation at 99°C in 2x Laemmli buffer (50mM 1,4-dithiothreitol; DTT) for 10 mins to elute all bound proteins. The released proteins were quantified by subjecting samples to a Bradford Protein assay and measuring absorbance at 590 nm with an Infinite M200 Pro plate reader (Tecan, Switzerland).

#### SDS-PAGE analysis

Protein samples were separated and visualized by denaturing SDS-PAGE. First, samples (0.1 mg/ml) were mixed at a 1:1 ratio with 2x Laemmli sample buffer containing 50 mM DTT and heated at 99°C for 10 min. They were then applied onto a 4-20% polyacrylamide gel (MiniPROTEAN TGX, BioRad) and run in SDS running buffer (25 mM Tris, 192 mM glycine, 1% (w/v) SDS, pH 8.3) at 200 V for 35 min. Gels were stained for proteins using Coomassie R-250 and subsequent gel images were captured on a Chemidoc MP Imaging System (Bio-Rad).

#### Liquid chromatography mass spectrometry (LC-MS)

For mass spectrometry analysis, the protein content of the recovered corona-NP complexes was quantified by Bicinchoninic acid (BCA) assay as per the manufacturer’s instructions. Normalized samples (0.1 mg/ml) were reduced and alkylated (HEPES - pH 8.5, 100 mM; sodium deoxycholate - SDC, 1%; tris(2-carboxyethyl)phosphine - 5 mM; 2- iodoacetamide - 10 mM) for 10 min at 95°C. For protein digestion, samples were incubated with sequencing grade trypsin (0.01 mg/mL; Promega Corporation) overnight at 37°C. Afterward, digestion reactions were halted by adding 10x volume of 90% acetonitrile (ACN) and 1% trifluoroacetic Acid (TFA), with any insoluble proteins removed by centrifugation. Soluble digested peptides were then loaded onto solid phase extraction (SPE) columns, equilibrated with 90% ACN and 1% TFA, by centrifugation at 2000 rcf for 2 min. Columns were then washed twice with 100 μL 10% ACN and 0.1% TFA, and the sample subsequently eluted with 50 μL of elution buffer (1M NH_4_OH_3_, 80% ACN). The resulting peptide-containing eluate was then collected and dried by speed vacuum for 2 h. Afterward, peptides were resuspended in 2% ACN and 0.2% TFA in water, with 1 μL samples loaded into a Q Exactive Plus Hybrid Quadrupole-Orbitrap Mass Spectrometer (ThermoFisher) *via* Acclaim PepMap 100 C18 LC columns (ThermoFisher). Samples analysis was performed using PEAKS Studio 8.5 software (Bioinformatics Solutions Inc.) and MASCOT (Matrix Science, UK) software. Only proteins that had been identified *via* at least one unique peptide were included. To account for variations in protein size, spectral counts were normalized by the molecular weight of each identified protein. Subsequently, the relative abundance of each protein within the evaluated protein corona was calculated using the normalized spectral counts.^[46, 104]^

#### Identified protein analysis

Proteins were categorized by molecular weight as determined by LC-MS output. Proteins were further categorized by physiological function as determined by PANTHER classification^[105]^ (or UniProt if not identified in PANTHER); net charge (at pH 7.4) and amino acid content as calculated by Prot Pi; and grand average of hydropathicity (GRAVY) was calculated using ExPASy ProtParam.

#### Protein enrichment/Depletion analysis

Enrichment and depletion was determined by calculating the Log_2_ of the fold change in RPA between any given protein identified in the protein corona and its corresponding source fluid:

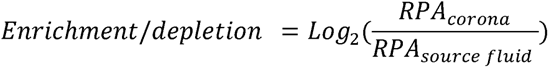

In this instance, the total corona was used to calculate RPA, as proteins detected within the top 200 of the coronas were not always detected within the top 200 of the source fluids. If a protein detected in the corona was not detected in the corresponding native fluid, then RPA was set to the lowest value detected in the fluid.

### Cellular interaction studies

#### Cell culture

For *in vitro* studies, bEnd.3 mouse brain endothelial cells; BV2 mouse microglial cells; and 1321N1 human astrocytoma cells were each cultured in Dulbecco’s Modified Eagle’s Medium (DMEM) supplemented with 10% fetal bovine serum (FBS) and 1% penicillin-streptomycin at 37°C in a 5% CO_2_-containing humidified incubator. SHSY-5Y human neuroblastoma cells were cultured under identical conditions, with the addition of Ham’s F-12 Nutrient Mix to the media (DMEM/F-12). Moreover, SHSY-5Y cells were differentiated into neurons using a method adapted from de Medeiros *et al.* ^[106]^ Here, the cells were initially seeded into either 6- or 96-well plates and incubated overnight (Day-1). The following day, culture media was replaced with DMEM/F-12 supplemented with 1% FBS. Subsequently, on Day-3, this media was replaced with fresh DMEM/F-12 media enriched with 1% FBS and 10 μM retinoic acid, which was repeated on Day-5 and Day-7. We have previously demonstrated several markers of neuronal maturation (i.e. development of characteristic polarized morphologies, extensive neurite arborization and up-regulation of the neuronal marker NeuN), as well as ∼50% higher activity of the enzyme acetylcholinesterase (a marker of mature cholinergic neurons), at day 7-10 *in vitro* using this protocol.^[107]^ Exemplar images showing SHSY-5Y differentiation into neurons are available in **Supplementary Figure S1**.

#### Flow cytometry

To simultaneously assess cell viability and nanoparticle internalization (with and without coronas) in various brain cell types, we employed flow cytometric analysis. Cells were seeded at a density of 5.0 x 10^3^ cells per well in 24-well culture plates and cultured at 37 °C for 24 h. Next, cells were incubated with 500 μg/mL of FITC-loaded nanoparticles (with and without coronas) in serum-free media (or media containing 1% FBS in the case of differentiated neurons) for 24 h. Next, supernatant from the treated cells was collected and stored for downstream cytokine analysis (see below). Cells were then washed with PBS to remove non-internalized particles, collected by trypsinization and resuspended in 0.5 mL of FACS buffer (PBS, 5 mM EDTA, 5% FBS), spiked with DAPI (1 µg/ml) and analyzed on the BD LSRII Flow Cytometer (BD Biosciences). Cells were initially identified using forward versus side scatter (FSC/SSC) gating, excluding cellular debris. The percentage of live single cells (i.e., cell viability) was determined by gating DAPI fluorescence (358 nm filter)/FSC. Uptake of FITC-loaded nanoparticles by live single cells was subsequently determined by further gating both DAPI/FITC fluorescence (488 nm filter). A minimum of 20,000 events were recorded for all samples assessed, with post-processing data analyzed using FlowJo software (Version 10.9, FlowJo LLC)

#### Fluorescent microscopy

The cellular uptake of FITC-loaded nanoparticles (with and without coronas) was also visualized by confocal microscopy. Here, cells were seeded on coverslips in a 6-well plate (1.0 x 10^4^ cells per well) for 24 h to allow cell adherence. Cells were then exposed to 500 μg/mL of FITC-loaded nanoparticles suspended in serum-free media and co-incubated for either 6 or 24 h. Thereafter, cells were washed with PBS three times to remove any non-internalized particles, and fixed with 4% paraformaldehyde for 15 min. After PBS wash, nuclei were stained with DAPI (2 μg/mL) in the dark for 20 min, and washed thrice with PBS. Finally, all cells were washed and the cover slips mounted onto glass slides for visualization using a Nikon A1R Laser scanning confocal inverted microscope and processed using the Nikon NIS Elements software.

#### Correlation analysis

Pearson’s correlation coefficients (*r)* were calculated to determine the relationship between uptake of NPs and the top 20 corona proteins using the GraphPad Prism software (v 9.1)

#### Cytokine assays

To evaluate immune responses to corona-nanoparticle complexes, we analyzed supernatants collected from the preceding flow cytometric experiments for the presence of pro-inflammatory cytokines TNF and IL-6. This analysis was performed using a cytokine bead array (CBA Mouse IL-6/TNF Flex Set, BD Biosciences) following the manufacturer’s instructions. Data was analyzed using BD CBA analysis software.

## ASSOCIATED CONTENT

### Supporting Information

Experimental studies investigating CSF protein coronas; Top 20 proteins in serum and CSF source fluids; Differentiation of SH-SY5Y neuroblastoma cells; confocal images showing uptake of fluorescent nanoparticles into various brain cell types; cell viability after co-incubation with increasing concentrations of nanoparticles with and without coronas.

### Notes

The authors declare no competing financial interest.

## ACKNOWLEDGMENTS

This work was supported by research grants awarded to A.C. and L.C.P. from: Dementia Australia Research Foundation; Mason Foundation; and the National Foundation for Medical Research and Innovation. A.C. is supported by a Chancellor’s Research Fellowship from the University of Technology Sydney (UTS). The authors acknowledge the technical and scientific assistance of both the UTS Proteomics, Lipidomics and Metabolomics Core Facility, and the UTS Microbial Imaging Facility. The authors also pay their respects to the Gadigal people, who are the traditional custodians of the land on which this research took place.

The authors would like to acknowledge Professor Anna Salvati (University of Groningen, Netherlands) for their help with the correlation analysis.

