## Supplementary information for "Protein Coronas Derived from Cerebrospinal Fluid Enhance the Interactions Between Nanoparticles and Brain Cells"

*Nabila Morshed^1^, Claire Rennie^1^, Matthew Faria^2^, Lyndsey Collins-Praino^3^, and Andrew Care^1^**

^1^School of Life Sciences, University of Technology Sydney, NSW 2007, Australia

^2^Department of Biomedical Engineering, The University of Melbourne, Victoria 3010, Australia

^3^School of Biomedicine, Faculty of Health and Medical Sciences, The University of Adelaide, SA 5005, Australia.

Contents

**Supplementary Table S1.** Experimental studies reporting CSF-derived protein coronas on nanoparticles.

**Supplementary Table S2.** Functional classification of Top-200 proteins identified in native fluids and nanoparticle-corona complexes.

**Supplementary Table S3.** Top-20 most abundant proteins identified in native rat serum and CSF.

**Supplementary Table S3.** Functional classifications of Top-200 proteins identified

**Supplementary Figure S1.** Differentiation of SH-SH5Y neuroblastoma cells into neurons.

**Supplementary Figure S3.** Uptake of FITC-loaded polymeric nanoparticles (with and without coronas) by different brain cell types.

**Supplementary Figure S4.** Viability of different brain cell types incubated with corona-nanoparticle complexes.

**Supplementary Table S1:** Summary of experimental studies reporting CSF-derived protein coronas on nanoparticles.

| **Nanoparticle type** | **Shape, Size, Charge** | **CSF source** | **Proteomic analysis** | **Biological**  **interactions studied** | **Notes** | **Ref** |
| --- | --- | --- | --- | --- | --- | --- |
| Polystyrene | Spherical  PEGylated: 117 nm  7 mV  Functionalized: 121  7 mV | Pooled rodent (Sprague Dawley Rats) CSF | LC/MS | Yes. Internalization into bEnd3 endothelial and C6 glioma cells | Effect on targeting antibodies | ^1^ |
| Polystyrene | Spherical 100 nm  -60 mV | Pooled normal human CSF | LC/MS | No | Focus on bio-nano interactions | ^2^ |
| Single-walled Carbon Nanotubes | Rod  100-1,000 (length) x 1(diameter) nm  -19 mV | Pooled normal human CSF | LC/MS | No | Focus on bio-nano interactions | ^2^ |
| Gold | Spherical 20 nm  32 mV | Human (lacks details) | 2D gel only | No | *In vitro* interactions of bio-corona with AB peptides | ^3^ |
| Gold | Rod  Aspect ratio ~ 3.5 nm  29 mV | Human (lacks details) | 2D gel only | No | *In vitro* interactions of bio-corona with AB peptides | ^3^ |
| Silica | Spherical  177 nm  -23 mV | Murine | No | Yes.  Internalization into BV2 cells | Effect of corona formation on SiO_2_ toxicity | ^4^ |

**Supplementary Table S2.** Functional classification of Top-200 proteins identified in native fluids and nanoparticle-corona complexes.

*Proteins classified as “Defense/immunity” and “Transmembrane signal receptor” were combined into the “Miscellaneous/unknown” classification due to low abundance*

| **Protein** | **Accession number** | **Functional classification** |
| --- | --- | --- |
| 11-beta-hydroxysteroid dehydrogenase type 2 | P50233\|DHI2_RAT | Metabolite interconversion enzyme |
| 14-3-3 protein beta/alpha | A0A8I6GMK2\|A0A8I6GMK2_RAT | Cytoskeletal protein/scaffold |
| 14-3-3 protein epsilon | A0A8I6GEP3\|A0A8I6GEP3_RAT | Cytoskeletal protein/scaffold |
| 14-3-3 protein gamma | P61983\|1433G_RAT | Cytoskeletal protein/scaffold |
| 14-3-3 protein theta | P68255\|1433T_RAT | Cytoskeletal protein/scaffold |
| 14-3-3 protein zeta/delta | P63102\|1433Z_RAT | Miscellaneous/unknown |
| 1-cysPrx_C domain-containing protein | A0A8I5Y2B8\|A0A8I5Y2B8_RAT | Metabolite interconversion enzyme |
| 2-hydroxyacylsphingosine 1-beta-galactosyltransferase 1 | Q09426\|CGT_RAT | Metabolite interconversion enzyme |
| Aa1018 | Q7TQ11\|Q7TQ11_RAT | Acute phase |
| Aa1249 | Q7TMA9\|Q7TMA9_RAT | Miscellaneous/unknown |
| Ac2-248 | Q7TPI9\|Q7TPI9_RAT | Protease Inhibitor |
| Actin related protein 2/3 complex, subunit 4 | A0A8I6APA3\|A0A8I6APA3_RAT | Cytoskeletal protein/scaffold |
| Actin, alpha cardiac muscle 1 | P68035\|ACTC_RAT\|A0A8I5ZVP6\| | Cytoskeletal protein/scaffold |
| Actin, alpha skeletal muscle | P68136\|ACTS_RAT | Cytoskeletal protein/scaffold |
| Actin, aortic smooth muscle | P62738\|ACTA_RAT | Cytoskeletal protein/scaffold |
| Actin, beta-like 2 | D3ZRN3\|D3ZRN3_RAT | Cytoskeletal protein/scaffold |
| Actin, cytoplasmic 1 | P60711\|ACTB_RAT\|A0A0G2K3K2\| | Cytoskeletal protein/scaffold |
| Actin, cytoplasmic 2 | P63259\|ACTG_RAT | Cytoskeletal protein/scaffold |
| Actin, gamma-enteric smooth muscle | P63269\|ACTH_RAT | Cytoskeletal protein/scaffold |
| Adenylate cyclase type 5 | Q04400\|ADCY5_RAT | Metabolite interconversion enzyme |
| Adenylyl cyclase-associated protein 1 | Q08163\|CAP1_RAT | Cytoskeletal protein/scaffold |
| Adenylyl cyclase-associated protein | A0A8I6G6M3\|A0A8I6G6M3_RAT | Cytoskeletal protein/scaffold |
| Adiponectin a | G3V7N9\|G3V7N9_RAT | Acute phase |
| Adiponectin b | A0A3B0IYY4\|A0A3B0IYY4_RAT | Acute phase |
| ADP-ribosylation factor 1 | P84079\|ARF1_RAT | Miscellaneous/unknown |
| ADP-ribosylation factor 2 | P84082\|ARF2_RAT | Miscellaneous/unknown |
| ADP-ribosylation factor 3 | P61206\|ARF3_RAT | Miscellaneous/unknown |
| ADP-ribosylation factor 5 | P84083\|ARF5_RAT | Miscellaneous/unknown |
| ADP-ribosylation factor | A0A8I5ZKF2\|A0A8I5ZKF2_RAT | Miscellaneous/unknown |
| Afamin | G3V9R9\|G3V9R9_RAT | Transfer/carrier protein/transporter |
| Albumin | A0A0G2JSH5\|A0A8I5Y4N3\|P02770\|ALBU_RAT | Acute phase |
| Alpha globin | Q63910\|Q63910_RAT | Transfer/carrier protein/transporter |
| Alpha-1-acid glycoprotein | A0A0H2UHF8\|A0A8I5ZRW3\|A0A8I5ZRW3_RAT | Acute phase |
| Alpha-1-antiproteinase | A0A0G2JY31\|A0A0G2JY31_RAT | Protease Inhibitor |
| Alpha-1B-glycoprotein | Q9EPH1\|A1BG_RAT | Immunoglobulins |
| Alpha-1-inhibitor 3 | A0A0G2K926\|P14046\|A1I3_RAT | Protease Inhibitor |
| Alpha-1-macroglobulin | Q63041\|A1M_RAT | Protease Inhibitor |
| Alpha-2-glycoprotein 1, zinc | Q3B8R6\|Q3B8R6_RAT | Defense/immunity protein |
| Alpha-2-HS-glycoprotein | A0A8I5Y586\|A0A8I5ZY55\|A0A8I5ZY55_RAT | Protease Inhibitor |
| Alpha-2-macroglobulin | P06238\|A2MG_RAT | Acute phase |
| Alpha-2u globulin | Q9JJH9\|Q9JJH9_RAT | Transfer/carrier protein/transporter |
| Alpha-amylase | Q99N59\|Q99N59_RAT | Miscellaneous/unknown |
| Alpha-enolase | P04764\|ENOA_RAT | Metabolite interconversion enzyme |
| Amidophosphoribosyltransferase | P35433\|PUR1_RAT | Metabolite interconversion enzyme |
| Amyloid-beta A4 protein | A0A8I5Y8G9\|A0A8I5Y8G9_RAT | Miscellaneous/unknown |
| Androgen-binding protein (Fragment) | D2XZ41\|D2XZ41_RAT | Miscellaneous/unknown |
| Angiotensin-converting enzyme | P47820\|ACE_RAT | Protein modifying enzyme |
| Angiotensinogen | P01015\|ANGT_RAT | Protease Inhibitor |
| Anionic trypsin-2 | P00763\|TRY2_RAT | Protein modifying enzyme |
| Annexin A2 | Q07936\|ANXA2_RAT | Miscellaneous/unknown |
| Annexin A4 | P55260\|ANXA4_RAT | Miscellaneous/unknown |
| Annexin A6 | P48037\|ANXA6_RAT | Miscellaneous/unknown |
| Annexin | A0A8J8XVA7\|F7FLB7\|F7FLB7_RAT | Miscellaneous/unknown |
| Antigen p97 (Melanoma associated) identified by monoclonal antibodies 133.2 and 96.5 (Predicted) | D4ADK7\|D4ADK7_RAT | Transfer/carrier protein/transporter |
| Antithrombin-III | Q5M7T5\|F7EY53\|F7EY53_RAT | Protease Inhibitor |
| Apolipoprotein A-I | A0A0G2JU23\|P04639\|APOA1_RAT | Apolipoprotein |
| Apolipoprotein A-II | P04638\|APOA2_RAT | Apolipoprotein |
| Apolipoprotein A-IV | A0A0G2JVX7\|P02651\|APOA4_RAT | Apolipoprotein |
| Apolipoprotein B-100 | A0A8I6AJ20\|Q7TMA5\|APOB_RAT | Apolipoprotein |
| Apolipoprotein C-I | A0A8I5ZRS9\|P19939\|APOC1_RAT | Apolipoprotein |
| Apolipoprotein C-II | G3V8D4\|APOC2_RAT | Apolipoprotein |
| Apolipoprotein C-III | A0A0H2UI39\|P06759\|APOC3_RAT | Apolipoprotein |
| Apolipoprotein C-IV | P55797\|APOC4_RAT | Apolipoprotein |
| Apolipoprotein D | P23593\|APOD_RAT | Apolipoprotein |
| Apolipoprotein E | A0A8I6ACI7\|A0A8I6ACI7\|P02650\|APOE_RAT | Apolipoprotein |
| Apolipoprotein M | A0A8I5YC12\|P14630\|APOM_RAT | Apolipoprotein |
| Apolipoprotein N | Q5M890\|Q5M890_RAT | Apolipoprotein |
| Aquaporin-1 | P29975\|AQP1_RAT | Transfer/carrier protein/transporter |
| Armadillo repeat-containing protein 5 | Q5PQP9\|ARMC5_RAT | Cytoskeletal protein/scaffold |
| ATP synthase subunit beta | G3V6D3\|G3V6D3_RAT | Transfer/carrier protein/transporter |
| Bardet-Biedl syndrome 9 | A0A1B0GWY0\|A0A1B0GWY0_RAT | Miscellaneous/unknown |
| Beta-2-glycoprotein 1 | Q5I0M1\|Q5I0M1_RAT | Complement component |
| Beta-2-microglobulin | P07151\|B2MG_RAT | Defense/immunity protein |
| Beta-enolase | P15429\|ENOB_RAT | Metabolite interconversion enzyme |
| Beta-hexosaminidase | A0A8I5ZT00\|A0A8I5ZT00_RAT | Miscellaneous/unknown |
| Biliverdin reductase B | A0A8I5ZUN1\|A0A8I5ZUN1_RAT | Metabolite interconversion enzyme |
| Brain acid soluble protein 1 | A0A8I6A304\|A0A8I6A304_RAT | Miscellaneous/unknown |
| BRCA1-associated ATM activator 1 | D3ZSM9\|D3ZSM9_RAT | Miscellaneous/unknown |
| Brevican core protein | G3V8G4\|G3V8G4_RAT | Extracellular matrix protein |
| Bromodomain and PHD finger-containing, 1 | D4A411\|D4A411_RAT | Miscellaneous/unknown |
| C3 and PZP-like, alpha-2-macroglobulin domain-containing 8 | A0A8I6AG69\|A0A8I5ZW14\|A0A8I6AMC9\|A0A8I6AMC9_RAT | Protease Inhibitor |
| C4b-binding protein alpha chain | Q5M891\|Q63514\|C4BPA_RAT | Complement component |
| C5a anaphylatoxin chemotactic receptor 1 | P97520\|C5AR1_RAT | Miscellaneous/unknown |
| Cadherin-1 | Q9R0T4\|CADH1_RAT | Miscellaneous/unknown |
| Cadherin-5 | F1M7E5\|F1M7E5_RAT | Miscellaneous/unknown |
| Calcium-transporting ATPase | A0A0G2K9Q6\|A0A0G2K9Q6_RAT | Transfer/carrier protein/transporter |
| Calmodulin 2 | D4ABV5\|D4ABV5_RAT | Miscellaneous/unknown |
| Calmodulin | A0A8I6GKN5\|A0A8I6GKN5_RAT | Miscellaneous/unknown |
| Calmodulin-1 | P0DP29\|CALM1_RAT | Miscellaneous/unknown |
| Calpain 11 | Q4V8Q1\|CAN11_RAT | Metabolite interconversion enzyme |
| Calsyntenin-1 | Q6Q0N0\|CSTN1_RAT | Miscellaneous/unknown |
| Carbonic anhydrase 2 | P27139\|CAH2_RAT | Metabolite interconversion enzyme |
| Carbonic anhydrase | A0A8I6AMP9\|A0A8I6AMP9_RAT | Metabolite interconversion enzyme |
| Carbonyl reductase [NADPH] 3 | B2GV72\|CRB3_RAT | Metabolite interconversion enzyme |
| Carboxylesterase 1C | P10959\|EST1C_RAT | Metabolite interconversion enzyme |
| Carboxylesterase 1E | Q63108\|EST1E_RAT | Metabolite interconversion enzyme |
| Carboxylic ester hydrolase | D3ZGK7\|D3ZGK7_RAT | Metabolite interconversion enzyme |
| Carboxypeptidase B2 | Q9EQV9\|CBPB2_RAT | Protein modifying enzyme |
| Carboxypeptidase E | P15087\|CBPE_RAT | Protein modifying enzyme |
| Carboxypeptidase N catalytic chain | A0A8I5ZTU6\|A0A8I5ZTU6_RAT | Metabolite interconversion enzyme |
| Carboxypeptidase N subunit 2 | F1LQT4\|F1LQT4_RAT | Transmembrane signal receptor |
| Carboxypeptidase Q | Q6IRK9\|CBPQ_RAT | Protein modifying enzyme |
| Cationic trypsin-3 | P08426\|TRY3_RAT | Protein modifying enzyme |
| Cationic trypsinogen | G3V7Q8\|G3V7Q8_RAT | Metabolite interconversion enzyme |
| CD5 antigen-like | Q4KM75\|Q4KM75_RAT | Miscellaneous/unknown |
| CD9 antigen | A0A8I6ALV0\|A0A8I6ALV0_RAT | Cytoskeletal protein/scaffold |
| CDK-activating kinase assembly factor MAT1 | A0A8I6ATM4\|A0A8I6ATM4_RAT | Miscellaneous/unknown |
| Cell adhesion molecule 3 | Q1WIM3\|CADM3_RAT | Miscellaneous/unknown |
| Ceruloplasmin | G3V7K3\|P13635\|CERU_RAT | Acute phase |
| Cfh protein | Q5XJW6\|Q5XJW6_RAT | Complement component |
| Chloride intracellular channel protein 1 | Q6MG61\|CLIC1_RAT | Transfer/carrier protein/transporter |
| Chloride intracellular channel protein 6 | Q811Q2\|CLIC6_RAT | Transfer/carrier protein/transporter |
| Chloride intracellular channel protein | F1M9X4\|G3V8C4\|G3V8C4_RAT | Transfer/carrier protein/transporter |
| Choline transporter-like protein 2 | B4F795\|CTL2_RAT | Transfer/carrier protein/transporter |
| Class I histocompatibility antigen, Non-RT1.A alpha-1 chain | P15978\|HA11_RAT | Defense/immunity protein |
| Clusterin | A0A0G2K259\|P05371\|CLUS_RAT | Miscellaneous/unknown |
| Coagulation factor IX | P16296\|FA9_RAT | Protein modifying enzyme |
| Coagulation factor V | A0A0G2K3W2\|A0A0G2K3W2_RAT | Miscellaneous/unknown |
| Coagulation factor X | A0A0H2UHR6\|A0A0H2UHR6_RAT | Protein modifying enzyme |
| Coagulation factor XII | A0A0H2UI19\|A0A0H2UI19_RAT | Metabolite interconversion enzyme |
| Cochlin | B1H259\|B1H259_RAT | Extracellular matrix protein |
| Codanin 1 | D4A1U4\|D4A1U4_RAT | Miscellaneous/unknown |
| Cofilin 2 | M0RC65\|M0RC65_RAT | Cytoskeletal protein/scaffold |
| Cofilin-1 | A0A8I5ZT72\|A0A8I5ZT72_RAT | Cytoskeletal protein/scaffold |
| Complement C1q subcomponent subunit A | A0A3B0J380\|P31720\|C1QA_RAT | Cytoskeletal protein/scaffold |
| Complement C1q subcomponent subunit B | P31721\|C1QB_RAT | Cytoskeletal protein/scaffold |
| Complement C1q subcomponent subunit C | A0A8I5ZWV4\|P31722\|C1QC_RAT | Cytoskeletal protein/scaffold |
| Complement C1s subcomponent | A0A8I6A2W2\|A0A8I6A2W2_RAT | Protein modifying enzyme |
| Complement C2 | A0A8J8YKQ8\|A0A8J8YKQ8_RAT | Complement component |
| Complement C3 | M0RBJ7\|M0RBF1\|A0A8I6ATP7\|P01026\|CO3_RAT | Acute phase |
| Complement C4 | P08649\|CO4_RAT | Acute phase |
| Complement C5 | A0A8I5ZDN9\|P08650\|CO5_RAT | Protease Inhibitor |
| Complement C8 alpha chain | A0A8I6ACZ6\|A0A8I6ACZ6_RAT | Complement component |
| Complement component C6 | F1M7F7\|Q811M5\|CO6_RAT | Complement component |
| Complement component C8 | A0A8I5ZWX9\|A0A8I5ZWX9_RAT | Complement component |
| Complement component C9 | F7F389\|Q62930\|CO9_RAT | Complement component |
| Complement factor B | G3V615\|G3V615_RAT | Complement component |
| Complement factor D | G3V7H3\|G3V7H3_RAT | Protein modifying enzyme |
| Complement factor H | G3V9R2\|G3V9R2_RAT | Complement component |
| Complement factor H-related 2 | A0A8I6ATZ0\|A0A8I6ATZ0_RAT | Complement component |
| Complement factor I | A0A8I6AC86\|A0A8I6AC86_RAT | Protein modifying enzyme |
| Complement factor properdin | B0BNN4\|B0BNN4_RAT | Complement component |
| Complement subcomponent C1r | B5DEH7\|B5DEH7_RAT | Protein modifying enzyme |
| Contactin-2 | G3V758\|G3V758_RAT | Miscellaneous/unknown |
| Corticosteroid-binding globulin | P31211\|CBG_RAT | Protease Inhibitor |
| C-reactive protein | P48199\|CRP_RAT | Acute phase |
| Creatine kinase B-type | P07335\|KCRB_RAT | Metabolite interconversion enzyme |
| Creatine kinase M-type | P00564\|KCRM_RAT | Metabolite interconversion enzyme |
| C-type lectin domain family 3, member B | D3ZUU6\|D3ZUU6_RAT | Miscellaneous/unknown |
| CXC chemokine RTCK1 | Q99ME0\|Q99ME0_RAT | Miscellaneous/unknown |
| Cystatin-C | A0A8I6AD91\|P14841\|CYTC_RAT | Protease Inhibitor |
| Dimethylaniline monooxygenase [N-oxide-forming] 4 | Q8K4B7\|FMO4_RAT | Metabolite interconversion enzyme |
| DnaJ (Hsp40) homolog, subfamily B, member 4 | Q5XIP0\|Q5XIP0_RAT | Miscellaneous/unknown |
| Ectonucleotide pyrophosphatase/phosphodiesterase family member 2 | A0A8I6ADH1\|A0A8I6ADH1_RAT | Metabolite interconversion enzyme |
| ELMO domain-containing 2 | D3ZNV6\|D3ZNV6_RAT | Cytoskeletal protein/scaffold |
| Elongation factor 1-alpha 1 | P62630\|EF1A1_RAT | Miscellaneous/unknown |
| Elongation factor 1-alpha 2 | P62632\|EF1A2_RAT | Miscellaneous/unknown |
| Elongation factor 1-alpha | M0R757\|M0R757_RAT | Miscellaneous/unknown |
| Epididymal-specific lipocalin-12 | A0A8I6A1Y6\|A0A8I6A1Y6_RAT | Transfer/carrier protein/transporter |
| Extracellular superoxide dismutase [Cu-Zn] | Q08420\|SODE_RAT | Metabolite interconversion enzyme |
| Ezrin | A0A8I6A8I6\|A0A8I6A8I6_RAT | Cytoskeletal protein/scaffold |
| Family with sequence similarity 120C | A0A1W2Q6F9\|A0A1W2Q6F9_RAT | Miscellaneous/unknown |
| Family with sequence similarity 178, member B | A0A8I6ABH8\|A0A8I6ABH8_RAT | Miscellaneous/unknown |
| Fc receptor-like 1 | A0A8I5ZXZ6\|A0A8I5ZXZ6_RAT | Miscellaneous/unknown |
| FERM domain-containing kindlin 3 | B2GVB9\|B2GVB9_RAT | Miscellaneous/unknown |
| FERM domain-containing protein | A0A8I5ZWW5\|A0A8I5ZWW5_RAT | Cytoskeletal protein/scaffold |
| Fetub protein | Q6IRS6\|Q6IRS6_RAT | Protease Inhibitor |
| Fetuin-B | A0A8I6AWQ8\|Q9QX79\|FETUB_RAT | Protease Inhibitor |
| Fibrinogen alpha chain | A0A8I6A5L4\|A0A8I6A5L4_RAT | Acute phase |
| Fibrinogen beta chain | A0A8I6A9U0\|A0A8I6A9U0_RAT | Acute phase |
| Fibrinogen gamma chain | P02680\|FIBG_RAT | Acute phase |
| Fibrinogen-like 2 | G3V7P2\|G3V7P2_RAT | Acute phase |
| Fibronectin | A0A8I6A5M1\|P04937\|FINC_RAT | Extracellular matrix protein |
| Ficolin (Collagen/fibrinogen domain containing) 1 | Q5M8B4\|Q5M8B4_RAT | Miscellaneous/unknown |
| Ficolin A | A0A8I6AG88\|A0A8I6AG88_RAT | Miscellaneous/unknown |
| Ficolin-1 | Q9WTS8\|FCN1_RAT | Miscellaneous/unknown |
| Filamin A | C0JPT7\|C0JPT7_RAT | Cytoskeletal protein/scaffold |
| Flavin-containing monooxygenase | A0A8I6AMY0\|A0A8I6AMY0_RAT | Metabolite interconversion enzyme |
| Flotillin | A0A0G2JU52\|A0A0G2JU52_RAT | Miscellaneous/unknown |
| Follistatin-related protein 1 | Q62632\|FSTL1_RAT | Protease Inhibitor |
| Fructose-bisphosphate aldolase A | P05065\|ALDOA_RAT | Metabolite interconversion enzyme |
| Fructose-bisphosphate aldolase | A0A8I6AKC0\|A0A8I6AKC0_RAT | Metabolite interconversion enzyme |
| Gamma-aminobutyric acid receptor subunit beta-1 | P15431\|GBRB1_RAT | Transfer/carrier protein/transporter |
| Gelsolin | A0A8I6A6D2\|Q68FP1\|GELS_RAT | Cytoskeletal protein/scaffold |
| Globin a4 | A0A1K0H3R5\|A0A1K0H3R5_RAT | Transfer/carrier protein/transporter |
| Glucose-6-phosphate isomerase | A0A8I6G6Z4\|A0A8I6G6Z4_RAT | Metabolite interconversion enzyme |
| Glutathione peroxidase 3 | P23764\|GPX3_RAT | Metabolite interconversion enzyme |
| Glutathione peroxidase 6 | Q64625\|GPX6_RAT | Metabolite interconversion enzyme |
| Glutathione S-transferase alpha-1 | P00502\|GSTA1_RAT | Metabolite interconversion enzyme |
| Glutathione S-transferase alpha-2 | P04903\|GSTA2_RAT | Metabolite interconversion enzyme |
| Glutathione S-transferase alpha-3 | P04904\|GSTA3_RAT | Metabolite interconversion enzyme |
| Glutathione S-transferase alpha-4 | P14942\|GSTA4_RAT | Transfer/carrier protein/transporter |
| Glutathione S-transferase alpha-5 | P46418\|GSTA5_RAT | Metabolite interconversion enzyme |
| Glutathione S-transferase Mu 1 | P04905\|GSTM1_RAT | Metabolite interconversion enzyme |
| Glutathione S-transferase Mu 2 | P08010\|GSTM2_RAT | Metabolite interconversion enzyme |
| Glutathione S-transferase Mu 5 | Q9Z1B2\|GSTM5_RAT | Metabolite interconversion enzyme |
| Glutathione S-transferase | A0A0G2K8Q5\|A0A0G2K8Q5_RAT | Metabolite interconversion enzyme |
| Glutathione S-transferase P | P04906\|GSTP1_RAT | Metabolite interconversion enzyme |
| Glutathione S-transferase, theta 3 | D3Z8I7\|D3Z8I7_RAT | Metabolite interconversion enzyme |
| Glyceraldehyde-3-phosphate dehydrogenase | P04797\|G3P_RAT | Metabolite interconversion enzyme |
| GM2 ganglioside activator | A0A8I6A5G9\|A0A8I6A5G9_RAT | Transfer/carrier protein/transporter |
| Gp_dh_C domain-containing protein | A0A8I5ZWK4\|F1LUI2\|F1LUI2_RAT | Metabolite interconversion enzyme |
| Granzyme M | G3V726\|G3V726_RAT | Protein modifying enzyme |
| GST N-terminal domain-containing protein | A0A8I6AQP1\|A0A8I6AQP1_RAT | Miscellaneous/unknown |
| Guanine nucleotide-binding protein G(i) subunit alpha-1 | A0A8I5Y7W9\|A0A8I5Y7W9_RAT | Miscellaneous/unknown |
| H2J.A histone | A0A8L2QY15\|A0A8L2QY15_RAT | Miscellaneous/unknown |
| Haptoglobin | A0A8I5ZPF0\|P06866\|HPT_RAT | Acute phase |
| Heat shock cognate 71 kDa protein | P63018\|HSP7C_RAT | Miscellaneous/unknown |
| Heat shock cognate protein 70, pseudogene 1 | M0RCB1\|M0RCB1_RAT | Miscellaneous/unknown |
| Heat shock protein HSP 90-alpha | P82995\|HS90A_RAT | Miscellaneous/unknown |
| Heat shock protein HSP 90-beta | A0A8I6AKS6\|A0A8I6AKS6_RAT | Miscellaneous/unknown |
| Heat shock-related 70 kDa protein 2 | P14659\|HSP72_RAT | Miscellaneous/unknown |
| HECT domain E3 ubiquitin protein ligase 2 | A0A0G2JYF3\|A0A0G2JYF3_RAT | Protein modifying enzyme |
| Hemoglobin alpha, adult chain 1 | A0A8I5ZYH2\|A0A0A0MP82\|A0A0A0MP82_RAT | Globin |
| Hemoglobin subunit alpha-1/2 | P01946\|HBA_RAT | Globin |
| Hemoglobin subunit beta-1 | A0A8I6AUL4\|P02091\|HBB1_RAT | Globin |
| Hemoglobin subunit beta-2 | P11517\|HBB2_RAT | Globin |
| Hemoglobin subunit zeta | G3V8R3\|G3V8R3_RAT | Globin |
| Hemopexin | P20059\|HEMO_RAT | Protein modifying enzyme |
| Heparin cofactor 2 | Q64268\|HEP2_RAT | Protease Inhibitor |
| Hermansky-Pudlak syndrome 5 protein homolog | A0A0G2JWL2\|Q7TMC3\|Q7TMC3_RAT | Miscellaneous/unknown |
| Heterogeneous nuclear ribonucleoprotein K | P61980\|HNRPK_RAT | Miscellaneous/unknown |
| Histidine-rich glycoprotein | A0A0G2K3G0\|Q99PS8\|HRG_RAT | Protease Inhibitor |
| Histone H2A | A0A8I6ASE8\|Q6I8Q6\|Q6I8Q6_RAT | Miscellaneous/unknown |
| Histone H2B | D3ZNH4\|G3V9C7\|G3V9C7_RAT | Miscellaneous/unknown |
| Histone H4 | P62804\|H4_RAT | Miscellaneous/unknown |
| IF rod domain-containing protein | A0A0G2JUG1\|A0A0G2JUG1_RAT | Cytoskeletal protein/scaffold |
| Ig delta chain C region (Fragment) | P01883\|IGHD_RAT | Immunoglobulins |
| Ig gamma-1 chain C region | P20759\|IGHG1_RAT | Immunoglobulins |
| Ig gamma-2A chain C region | P20760\|IGG2A_RAT | Immunoglobulins |
| Ig gamma-2B chain C region | P20761\|IGG2B_RAT | Immunoglobulins |
| Ig gamma-2C chain C region | P20762\|IGG2C_RAT | Immunoglobulins |
| Ig heavy chain V region IR2 | P01805\|HVR01_RAT | Immunoglobulins |
| Ig kappa chain C region, A allele | P01836\|KACA_RAT | Immunoglobulins |
| Ig kappa chain C region, B allele | P01835\|KACB_RAT | Immunoglobulins |
| Ig kappa chain V region S211 | P01681\|KVX01_RAT | Immunoglobulins |
| Ig lambda-2 chain C region | P20767\|LAC2_RAT | Immunoglobulins |
| Igh protein-like | D4ADK9\|D4ADK9_RAT | Immunoglobulins |
| IGv domain-containing protein | A0A8I5Y7P7\|A0A8I5Y7P7_RAT | Immunoglobulins |
| Immunoglobulin heavy constant epsilon | A0A0G2JYX2\|F1LN61\|F1LM30\|F1LM30_RAT | Immunoglobulins |
| Immunoglobulin joining chain | G3V6G1\|G3V6G1_RAT | Immunoglobulins |
| Importin subunit beta-1 | P52296\|IMB1_RAT | Transfer/carrier protein/transporter |
| Inducible T-cell co-stimulator ligand | A0A8I6ACR9\|A0A8I6ACR9_RAT | Immunoglobulins |
| Insulin-like growth factor II | P01346\|IGF2_RAT | Miscellaneous/unknown |
| Insulin-like growth factor-binding protein complex acid labile subunit | P35859\|ALS_RAT | Transmembrane signal receptor |
| Inter alpha-trypsin inhibitor, heavy chain 4 | Q5EBC0\|Q5EBC0_RAT | Protease Inhibitor |
| Inter-alpha trypsin inhibitor, heavy chain 1 | F7EYX4\|F7EYX4_RAT | Protease Inhibitor |
| Inter-alpha-trypsin inhibitor heavy chain 2 | A0A8I6A7A2\|D3ZFH5\|D3ZFH5_RAT | Protease Inhibitor |
| Inter-alpha-trypsin inhibitor heavy chain H3 | A0A8I5ZPG2\|A0A8I6ALV3\|A0A8I6GKX5\|A0A8I6A7A2\|Q63416\|ITIH3_RAT | Protease Inhibitor |
| Interleukin 31 | A0A8I5ZNP8\|A0A8I5ZNP8_RAT | Miscellaneous/unknown |
| Inward rectifier potassium channel 13 | O70617\|KCJ13_RAT | Transfer/carrier protein/transporter |
| Kallikrein B, plasma 1 | Q5FVS2\|Q5FVS2_RAT | Metabolite interconversion enzyme |
| Kelch-like 6 (Drosophila) (Predicted) | D4A1Z1\|D4A1Z1_RAT | Cytoskeletal protein/scaffold |
| Keratin 14 | Q6IFV0\|Q6IFV0_RAT | Cytoskeletal protein/scaffold |
| Keratin 16 | A0A0G2JXJ9\|Q6IFU9\|Q6IFU9_RAT | Cytoskeletal protein/scaffold |
| Keratin 71 | D3ZXB7\|D3ZXB7_RAT | Cytoskeletal protein/scaffold |
| Keratin 79 | A0A8I5ZR22\|A0A8I5ZR22_RAT | Cytoskeletal protein/scaffold |
| Keratin 82 | G3V939\|G3V939_RAT | Cytoskeletal protein/scaffold |
| Keratin, type I cytoskeletal 10 | Q6IFW6\|K1C10_RAT | Cytoskeletal protein/scaffold |
| Keratin, type I cytoskeletal 14 | Q6IFV1\|K1C14_RAT | Cytoskeletal protein/scaffold |
| Keratin, type II cytoskeletal 1 | A0A0G2JST3\|Q6IMF3\|K2C1_RAT | Cytoskeletal protein/scaffold |
| Keratin, type II cytoskeletal 5 | Q6P6Q2\|K2C5_RAT | Cytoskeletal protein/scaffold |
| Kininogen 1 | Q5PQU1\|Q5PQU1_RAT | Protease Inhibitor |
| Kininogen 2 | A0A0G2JVQ5\|A0A0G2JVQ5\|A0A8I5ZK39\|A0A8I5ZK39_RAT | Protease Inhibitor |
| Kininogen-1 | F7EUK4\|A0A0G2KA54\|P08934\|KNG1_RAT | Protease Inhibitor |
| Lactotransferrin | D3ZAB1\|D3ZAB1_RAT | Transfer/carrier protein/transporter |
| Lactoylglutathione lyase | Q6P7Q4\|LGUL_RAT | Metabolite interconversion enzyme |
| Ldh_1_N domain-containing protein | A0A8I6GK09\|A0A8I6GK09_RAT | Metabolite interconversion enzyme |
| Leukemia inhibitory factor receptor | A0A8I5ZRS1\|G3V7K2\|G3V7K2_RAT | Transmembrane signal receptor |
| Leukocyte elastase inhibitor A | Q4G075\|ILEUA_RAT | Protease Inhibitor |
| Lipocln_cytosolic_FA-bd_dom domain-containing protein | F1M6Y6\|F1M6Y6_RAT | Transfer/carrier protein/transporter |
| Lipopolysaccharide-binding protein | A0A8I6AMU5\|Q3MID7\|Q3MID7_RAT | Defense/immunity protein |
| Liver carboxylesterase 1F | Q64573\|EST1F_RAT | Metabolite interconversion enzyme |
| Liver carboxylesterase B-1 | Q63010\|EST5_RAT | Metabolite interconversion enzyme |
| L-lactate dehydrogenase A chain | P04642\|LDHA_RAT | Metabolite interconversion enzyme |
| L-lactate dehydrogenase B chain | P42123\|LDHB_RAT | Metabolite interconversion enzyme |
| L-lactate dehydrogenase C chain | H9N9H4\|H9N9H4_RAT | Metabolite interconversion enzyme |
| L-lactate dehydrogenase | A0A8I6AAB9\|A0A8I6AAB9_RAT | Metabolite interconversion enzyme |
| LRRGT00077 | Q6TUG7\|Q6TUG7_RAT | Miscellaneous/unknown |
| LRRGT00083 | Q6TUG1\|Q6TUG1_RAT | Miscellaneous/unknown |
| LUC7-like 3 pre-mRNA splicing factor | A0A8I5ZVZ0\|A0A8I5ZVZ0_RAT | Miscellaneous/unknown |
| Lumican | P51886\|LUM_RAT | Cytoskeletal protein/scaffold |
| Lysosome-associated membrane glycoprotein 2 | A0A8I6AHB3\|A0A8I6AHB3_RAT | Miscellaneous/unknown |
| Lysozyme C-1 | P00697\|LYSC1_RAT | Metabolite interconversion enzyme |
| Macrophage colony-stimulating factor 1 receptor | A0A8I6A5F0\|A0A8I6A5F0_RAT | Transmembrane signal receptor |
| Major facilitator superfamily domain containing 8 | A0A8I6ANA0\|A0A8I6ANA0_RAT | Transfer/carrier protein/transporter |
| Major urinary protein | P02761\|MUP_RAT | Transfer/carrier protein/transporter |
| Malate dehydrogenase | A0A8I6A721\|A0A8I6A721_RAT | Metabolite interconversion enzyme |
| Mediator of RNA polymerase II transcription subunit 6 | A0A8I6AVM6\|A0A8I6AVM6_RAT | Miscellaneous/unknown |
| Melanotransferrin | A0A8I6AJB3\|A0A8I6AJB3_RAT | Transfer/carrier protein/transporter |
| Membrane associated ring-CH-type finger 6 | A0A8I5ZPM5\|A0A8I5ZPM5_RAT | Protein modifying enzyme |
| Merlin | G3V717\|G3V717_RAT | Cytoskeletal protein/scaffold |
| Metalloproteinase inhibitor 3 | P48032\|TIMP3_RAT | Protease Inhibitor |
| Mimecan | A0A1W2Q6Q0\|A0A1W2Q6Q0_RAT | Miscellaneous/unknown |
| MNAT1 component of CDK activating kinase | A0A8I5ZV40\|A0A8I5ZV40_RAT | Metabolite interconversion enzyme |
| Moesin | A0A8I5ZUJ9\|A0A8I5ZUJ9_RAT | Cytoskeletal protein/scaffold |
| Multidrug and toxin extrusion protein | D4A4W2\|D4A4W2_RAT | Transfer/carrier protein/transporter |
| Murinoglobulin-2 | A0A0G2JUW7\|A0A8I6AQK4\|Q6IE52\|MUG2_RAT | Protease Inhibitor |
| Myl6 protein | B2GV99\|B2GV99_RAT | Cytoskeletal protein/scaffold |
| Myoglobin | A0A8I6AHY4\|A0A8I6AHY4_RAT | Globin |
| Myosin light polypeptide 6 | A0A0G2JWE1\|Q64119\|MYL6_RAT | Cytoskeletal protein/scaffold |
| Myosin-9 | Q62812\|MYH9_RAT | Cytoskeletal protein/scaffold |
| Na(+)/H(+) exchange regulatory cofactor NHE-RF1 | Q9JJ19\|NHRF1_RAT | Cytoskeletal protein/scaffold |
| NACHT, LRR and PYD domains-containing protein 3 | D4A523\|NLRP3_RAT | Defense/immunity protein |
| NEL-like 2 (Chicken) | Q561K2\|Q561K2_RAT | Miscellaneous/unknown |
| Neural cell adhesion molecule 1 | A0A8I5ZQ32\|A0A8I5ZQ32_RAT | Miscellaneous/unknown |
| Neutrophilic granule protein | D3ZY96\|D3ZY96_RAT | Defense/immunity protein |
| NFKB inhibitor zeta | D4A1S9\|D4A1S9_RAT | Miscellaneous/unknown |
| NLR family, pyrin domain-containing 14 | F1M918\|F1M918_RAT | Defense/immunity protein |
| Olfactory receptor | A0A8I6A622\|A0A8I6A622_RAT | Miscellaneous/unknown |
| Pannexin-3 | P60572\|PANX3_RAT | Transfer/carrier protein/transporter |
| Paraoxonase | A0A8I5ZN72\|A0A8I5ZN72_RAT | Metabolite interconversion enzyme |
| Parvalbumin alpha | P02625\|PRVA_RAT | Miscellaneous/unknown |
| Peptidoglycan recognition protein 2 | M0R485\|M0R485_RAT | Defense/immunity protein |
| Peptidyl-prolyl cis-trans isomerase A | P10111\|PPIA_RAT | Miscellaneous/unknown |
| Peptidyl-prolyl cis-trans isomerase B | P24368\|PPIB_RAT | Miscellaneous/unknown |
| Peptidyl-prolyl cis-trans isomerase | A0A8I6A629\|Q6AYQ9\|Q6AYQ9_RAT | Miscellaneous/unknown |
| Peroxiredoxin-1 | Q63716\|PRDX1_RAT | Metabolite interconversion enzyme |
| Peroxiredoxin-2 | A0A8I6G2K0\|A0A8I6G2K0_RAT | Metabolite interconversion enzyme |
| Peroxiredoxin-5 | A0A8I5ZR89\|A0A8I5ZR89_RAT | Metabolite interconversion enzyme |
| Peroxiredoxin-6 | A0A8I6A038\|A0A8I6A038_RAT | Metabolite interconversion enzyme |
| Phosphatidylcholine-sterol acyltransferase | P18424\|LCAT_RAT | Metabolite interconversion enzyme |
| Phosphatidylethanolamine-binding protein 1 | A0A8I5ZQN0\|A0A8I5ZQN0_RAT | Protease Inhibitor |
| Phosphatidylinositol-glycan-specific phospholipase D | G3V8B1\|G3V8B1_RAT | Metabolite interconversion enzyme |
| Phosphoglucomutase-1 | A0A8I6AV00\|A0A8I6AV00_RAT | Metabolite interconversion enzyme |
| Phosphoglycerate kinase 1 | P16617\|PGK1_RAT | Metabolite interconversion enzyme |
| Phosphoglycerate kinase | A0A096MJL6\|A0A096MJL6_RAT | Metabolite interconversion enzyme |
| Phosphoglycerate mutase 1 | P25113\|PGAM1_RAT | Metabolite interconversion enzyme |
| Phosphoglycerate mutase 2 | P16290\|PGAM2_RAT | Metabolite interconversion enzyme |
| Phospholemman | O08589\|PLM_RAT | Transfer/carrier protein/transporter |
| Phospholipid transfer protein | A0A8I6AGE7\|E9PSP1\|E9PSP1_RAT | Defense/immunity protein |
| Phosphopyruvate hydratase | A0A8I6AR18\|M0R5J4\|M0R5J4_RAT | Metabolite interconversion enzyme |
| Plasma kallikrein 1 | P14272\|KLKB1_RAT | Protein modifying enzyme |
| Plasma membrane calcium-transporting ATPase 2 | P11506\|AT2B2_RAT | Transfer/carrier protein/transporter |
| Plasma membrane calcium-transporting ATPase 3 | Q64568\|AT2B3_RAT | Transfer/carrier protein/transporter |
| Plasma protease C1 inhibitor | A0A8I6ABI1\|A0A8I6A6T2\|Q6P734\|IC1_RAT | Protease Inhibitor |
| Plasminogen | A0A8I5ZVK5\|Q01177\|PLMN_RAT | Protein modifying enzyme |
| Platelet factor 4 | P06765\|PLF4_RAT | Miscellaneous/unknown |
| Platelet-activating factor acetylhydrolase | Q5M7T7\|Q5M7T7_RAT | Protein modifying enzyme |
| Polyubiquitin | A0A8I6A5Y3\|A0A8I6A5Y3_RAT | Miscellaneous/unknown |
| Potassium-transporting ATPase alpha chain 1 | P09626\|ATP4A_RAT | Transfer/carrier protein/transporter |
| Pregnancy-zone protein | A0A8L2Q3W7\|A0A8I6A2U4\|A0A8I6A2U4_RAT | Protease Inhibitor |
| Profilin | A0A8I6AKB2\|A0A8I6AKB2_RAT | Cytoskeletal protein/scaffold |
| Profilin-1 | P62963\|PROF1_RAT | Cytoskeletal protein/scaffold |
| Programmed cell death 5 | D4ADF5\|D4ADF5_RAT | Miscellaneous/unknown |
| Prohibitin | A0A8I5ZUY9\|A0A8I5ZUY9_RAT | Miscellaneous/unknown |
| Prosaposin | Q6P7A4\|Q6P7A4_RAT | Cytoskeletal protein/scaffold |
| Prostaglandin-H2 D-isomerase | P22057\|PTGDS_RAT | Transfer/carrier protein/transporter |
| Prostate stem cell antigen | A0A8I6A8F0\|A0A8I6A8F0_RAT | Miscellaneous/unknown |
| Protein AMBP | Q64240\|AMBP_RAT | Protease Inhibitor |
| Protein disulfide-isomerase A3 | P11598\|PDIA3_RAT | Miscellaneous/unknown |
| Protein disulfide-isomerase | A0A0H2UHM5\|A0A0H2UHM5_RAT | Miscellaneous/unknown |
| Protein FAM3C | A0A0G2JUB0\|A0A0G2JUB0_RAT | Defense/immunity protein |
| Protein kinase C-binding protein NELL2 | Q62918\|NELL2_RAT | Miscellaneous/unknown |
| Protein PHTF1 | F1M8G0\|PHTF1_RAT | Miscellaneous/unknown |
| Protein S100-A8 | P50115\|S10A8_RAT | Miscellaneous/unknown |
| Protein S100-A9 | A0A0H2UHJ1\|P50116\|S10A9_RAT | Miscellaneous/unknown |
| Protein Z, vitamin K-dependent plasma glycoprotein | A0A8I6A2L4\|A0A8I6A2L4_RAT | Metabolite interconversion enzyme |
| Protein Z-dependent protease inhibitor | Q62975\|ZPI_RAT | Protease Inhibitor |
| Prothrombin | P18292\|THRB_RAT | Protein modifying enzyme |
| Purine nucleoside phosphorylase | P85973\|PNPH_RAT | Metabolite interconversion enzyme |
| Putative gustatory receptor clone PTE03 (Fragment) | P35895\|GU03_RAT | Transmembrane signal receptor |
| Putative lysozyme C-2 | Q05820\|LYSC2_RAT | Metabolite interconversion enzyme |
| Pyruvate dehydrogenase phosphatase regulatory subunit | A0A8I5ZQQ3\|A0A8I5ZQQ3_RAT | Metabolite interconversion enzyme |
| Pyruvate kinase | M0RD14\|M0RD14_RAT | Metabolite interconversion enzyme |
| Pyruvate kinase PKM | P11980\|KPYM_RAT | Metabolite interconversion enzyme |
| RAB10, member RAS oncogene family | Q5RKJ9\|Q5RKJ9_RAT | Miscellaneous/unknown |
| RAB1A, member RAS oncogene family | E9PU16\|E9PU16_RAT | Miscellaneous/unknown |
| RAB1B, member RAS oncogene family | G3V6H0\|G3V6H0_RAT | Miscellaneous/unknown |
| Radixin | A0A8I5ZR70\|A0A8I5ZR70_RAT | Cytoskeletal protein/scaffold |
| Ras-related C3 botulinum toxin substrate 1 | Q6RUV5\|RAC1_RAT | Protease Inhibitor |
| Ras-related protein Rab-11A | P62494\|RB11A_RAT | Miscellaneous/unknown |
| Ras-related protein Rab-11B | A0A8I5ZWV2\|O35509\|RB11B_RAT | Miscellaneous/unknown |
| Ras-related protein Rab-12 | A0A8I5Y4W3\|A0A8I5Y4W3_RAT | Miscellaneous/unknown |
| Ras-related protein Rab-1A | Q6NYB7\|RAB1A_RAT | Miscellaneous/unknown |
| Ras-related protein Rab-1B | P10536\|RAB1B_RAT | Miscellaneous/unknown |
| Ras-related protein Rab-6A | Q9WVB1\|RAB6A_RAT | Miscellaneous/unknown |
| Ras-related protein Rap-1A | P62836\|RAP1A_RAT | Miscellaneous/unknown |
| Ras-related protein Rap-1b | Q62636\|RAP1B_RAT | Miscellaneous/unknown |
| RCG20603 | A0A0G2JSK1\|A0A0G2JSK1_RAT | Protease Inhibitor |
| RCG28243 | D3ZQV0\|D3ZQV0_RAT | Protein modifying enzyme |
| RCG31390 | G3V9A3\|G3V9A3_RAT | Cytoskeletal protein/scaffold |
| RCG32020 | D3ZV67\|D3ZV67_RAT | Miscellaneous/unknown |
| RCG41069 | A0A0G2JVF1\|A0A0G2JVF1_RAT | Miscellaneous/unknown |
| RCG43931 | Q6AYF8\|Q6AYF8_RAT | Protease Inhibitor |
| RCG44069 | Q68FX2\|Q68FX2_RAT | Protease Inhibitor |
| RCG50143 | A0A8I6AN99\|A0A8I6AN99_RAT | Miscellaneous/unknown |
| RCG53372 | A0A8I6AK42\|A0A8I6AK42_RAT | Immunoglobulins |
| Receptor for retinol uptake STRA6 | Q4QR83\|STRA6_RAT | Transfer/carrier protein/transporter |
| Receptor protein-tyrosine kinase | A0A8I5ZZ98\|A0A8I5ZZ98_RAT | Transmembrane signal receptor |
| Regulator of G-protein signaling 22 | A0A8I6AM59\|A0A8I6AM59_RAT | Miscellaneous/unknown |
| Regulator of microtubule dynamics protein 1 | A0A0G2K167\|A0A0G2K167_RAT | Cytoskeletal protein/scaffold |
| Retinol-binding protein 1 | P02696\|RET1_RAT | Acute phase |
| Retinol-binding protein 4 | P04916\|RET4_RAT | Acute phase |
| RGD1561426 | A0A8I5ZSU3\|A0A8I5ZSU3_RAT | Miscellaneous/unknown |
| Rho GDP-dissociation inhibitor 1 | Q5XI73\|GDIR1_RAT | Miscellaneous/unknown |
| RUN domain containing 3A | A0A0G2JW67\|A0A0G2JW67_RAT | Miscellaneous/unknown |
| Secretoglobin, family 2B, member 2 | F1LSA5\|F1LSA5_RAT | Miscellaneous/unknown |
| Selenoprotein P | A0A0G2JU99\|A0A0G2JU99_RAT | Metabolite interconversion enzyme |
| Serine (or cysteine) peptidase inhibitor, clade B, member 1b | F8WGA3\|F8WGA3_RAT | Protease Inhibitor |
| Serine (Or cysteine) proteinase inhibitor, clade A (Alpha-1 antiproteinase, antitrypsin), member 4 | Q5M8C3\|Q5M8C3_RAT | Protease Inhibitor |
| Serine protease 1 | P00762\|TRY1_RAT | Protein modifying enzyme |
| Serine protease inhibitor 2.1 (Fragment) | P09005\|SPI21_RAT | Protease Inhibitor |
| Serine protease inhibitor A3K | P05545\|SPA3K_RAT | Protease Inhibitor |
| Serine protease inhibitor A3L | P05544\|SPA3L_RAT | Protease Inhibitor |
| Serine protease inhibitor A3M | F1LR92\|F1LR92_RAT | Protease Inhibitor |
| Serine protease inhibitor A3N | A0A0G2KB85\|A0A0H2UHI5\|P09006\|SPA3N_RAT | Protease Inhibitor |
| Serotransferrin | P12346\|TRFE_RAT | Transfer/carrier protein/transporter |
| SERPIN domain-containing protein | A0A8I5ZRU6\|A0A8I5ZRU6_RAT | Protease Inhibitor |
| Serpin family A member 4 | A0A8I5ZZZ2\|A0A8I5ZZZ2_RAT | Protease Inhibitor |
| Serpin family A member 5 | F7EMJ6\|F7EMJ6_RAT | Protease Inhibitor |
| Serpin family B member 6A | A0A8I6A6H8\|A0A8I6A6H8_RAT | Protease Inhibitor |
| Serpin family F member 1 | A0A096MIW4\|A0A8J8XL70\|A0A8J8XL70_RAT | Protease Inhibitor |
| Serpin family F member 2 | F7FHF3\|A0A8I6AQI7\|A0A8I6AQI7_RAT | Protease Inhibitor |
| Serum amyloid P-component | A0A0H2UHH2\|A0A0H2UHH2_RAT | Miscellaneous/unknown |
| Serum paraoxonase/arylesterase 1 | P55159\|PON1_RAT | Metabolite interconversion enzyme |
| Short chain dehydrogenase/reductase family 42E, member 2 | A0A8I6A8Y0\|A0A8I6A8Y0_RAT | Metabolite interconversion enzyme |
| Signal peptidase complex catalytic subunit SEC11 | A0A8I5XVF9\|A0A8I5XVF9_RAT | Metabolite interconversion enzyme |
| Signal peptidase complex catalytic subunit SEC11A | P42667\|SC11A_RAT | Protein modifying enzyme |
| Signal recognition particle receptor subunit beta | Q7TMC7\|Q7TMC7_RAT | Transfer/carrier protein/transporter |
| Similar to 14-3-3 protein sigma | Q5EBB0\|Q5EBB0_RAT | Cytoskeletal protein/scaffold |
| Similar to Actin, cytoplasmic 2 (Gamma-actin) | A0A8I5Y704\|A0A8I5Y704_RAT | Cytoskeletal protein/scaffold |
| Similar to alpha-2u globulin PGCL2 | A0A096MK41\|A0A096MK41_RAT | Miscellaneous/unknown |
| Similar to BC049975 protein | F1M8F5\|F1M8F5_RAT | Protease Inhibitor |
| Similar to glyceraldehyde-3-phosphate dehydrogenase (phosphorylating) | A0A0G2K8S2\|A0A0G2K8S2_RAT | Metabolite interconversion enzyme |
| Similar to heat shock protein 1, alpha | A0A8I5ZQP6\|A0A8I5ZQP6_RAT | Miscellaneous/unknown |
| Similar to heat shock protein 8 | A0A8I6AQL9\|A0A8I6AQL9_RAT | Miscellaneous/unknown |
| Similar to Ig variable region, light chain | D3ZYE2\|D3ZYE2_RAT | Immunoglobulins |
| Similar to immunoglobulin kappa-chain VK-1 | M0RCN6\|M0RCN6_RAT | Immunoglobulins |
| Similar to polyubiquitin | A0A8I6A1Y5\|A0A8I6A1Y5_RAT | Miscellaneous/unknown |
| Similar to RIKEN cDNA 1300017J02 | A0A0G2K896\|A0A0G2K896_RAT | Transfer/carrier protein/transporter |
| Similar to Vanin-3 (Predicted) | D4A183\|D4A183_RAT | Metabolite interconversion enzyme |
| Sodium/potassium-transporting ATPase subunit alpha | A0A8I5Y5B0\|G3V8S4\|G3V8S4_RAT | Transfer/carrier protein/transporter |
| Sodium/potassium-transporting ATPase subunit beta | A0A096MJI9\|A0A096MJI9_RAT | Transfer/carrier protein/transporter |
| Sodium/potassium-transporting ATPase subunit beta-1 | P07340\|AT1B1_RAT | Transfer/carrier protein/transporter |
| Solute carrier family 12 member 2 | E9PTX9\|E9PTX9_RAT | Transfer/carrier protein/transporter |
| Solute carrier organic anion transporter family member 1A4 | A0A8L2QP25\|A0A8L2QP25_RAT | Transfer/carrier protein/transporter |
| SPARC-like protein 1 | P24054\|SPRL1_RAT | Extracellular matrix protein |
| Stathmin | A0A0G2K8P5\|A0A0G2K8P5_RAT | Cytoskeletal protein/scaffold |
| Superoxide dismutase [Cu-Zn] | A0A8I6G9Z6\|A0A8I6G9Z6_RAT | Metabolite interconversion enzyme |
| Talin 1 | A0A8I6ABG6\|A0A8I6ABG6_RAT | Cytoskeletal protein/scaffold |
| Tenascin XB | A0A8I6AE34\|A0A8I6AE34_RAT | Cytoskeletal protein/scaffold |
| Testis-expressed 46 | D4A3Q6\|D4A3Q6_RAT | Miscellaneous/unknown |
| Tetraspanin | A0A8I5ZZU3\|A0A8I5ZZU3_RAT | Cytoskeletal protein/scaffold |
| Thioredoxin-dependent peroxiredoxin | A0A8I5ZXJ7\|A0A8I5ZXJ7_RAT | Metabolite interconversion enzyme |
| Thrombospondin 1 | A0A0G2JV24\|A0A0G2JV24_RAT | Miscellaneous/unknown |
| T-kininogen 1 | P01048\|KNT1_RAT | Protease Inhibitor |
| T-kininogen 2 | P08932\|KNT2_RAT | Protease Inhibitor |
| Transferrin receptor protein 1 | A0A8I6AS83\|Q99376\|TFR1_RAT | Protein modifying enzyme |
| Transgelin | A0A8I5ZRB4\|A0A8I5ZRB4_RAT | Cytoskeletal protein/scaffold |
| Transgelin-2 | Q5XFX0\|TAGL2_RAT | Cytoskeletal protein/scaffold |
| Transgelin-3 | P37805\|TAGL3_RAT | Cytoskeletal protein/scaffold |
| Transmembrane serine protease 12 | F1M348\|F1M348_RAT | Protein modifying enzyme |
| Transthyretin | P02767\|TTHY_RAT | Acute phase |
| Triosephosphate isomerase | A0A0G2JWU1\|A0A0G2JWU1_RAT | Metabolite interconversion enzyme |
| Tropomyosin 3 | A0A8I6B5K9\|A0A8I6B5K9_RAT | Cytoskeletal protein/scaffold |
| Tropomyosin alpha-3 chain | Q63610\|TPM3_RAT | Cytoskeletal protein/scaffold |
| Tropomyosin alpha-4 chain | A0A0G2K2G8\|A0A0G2K2G8_RAT | Cytoskeletal protein/scaffold |
| Tropomyosin beta chain | A0A8I5ZR83\|A0A8I5ZR83_RAT | Cytoskeletal protein/scaffold |
| Tr-type G domain-containing protein | A0A8I6AKD4\|F1M6C2\|F1M6C2_RAT | Miscellaneous/unknown |
| Tubulin alpha chain | A0A8I6ALV8\|A0A8I6ALV8_RAT | Cytoskeletal protein/scaffold |
| Tubulin alpha-1A chain | P68370\|TBA1A_RAT | Cytoskeletal protein/scaffold |
| Tubulin alpha-1B chain | Q6P9V9\|TBA1B_RAT | Cytoskeletal protein/scaffold |
| Tubulin alpha-1C chain | Q6AYZ1\|TBA1C_RAT | Cytoskeletal protein/scaffold |
| Tubulin alpha-3 chain | Q68FR8\|TBA3_RAT | Cytoskeletal protein/scaffold |
| Tubulin alpha-4A chain | Q5XIF6\|TBA4A_RAT | Cytoskeletal protein/scaffold |
| Tubulin beta chain | M0R8B6\|M0R8B6_RAT | Cytoskeletal protein/scaffold |
| Tubulin beta-2A chain | A0A8L2UJU7\|P85108\|TBB2A_RAT | Cytoskeletal protein/scaffold |
| Tubulin beta-2B chain | Q3KRE8\|TBB2B_RAT | Cytoskeletal protein/scaffold |
| Tubulin beta-4B chain | Q6P9T8\|TBB4B_RAT | Cytoskeletal protein/scaffold |
| Tubulin beta-5 chain | A0A8L2RA69\|P69897\|TBB5_RAT | Cytoskeletal protein/scaffold |
| Type II keratin Kb15 | A0A0G2JVA8\|A0A0G2JVA8_RAT | Cytoskeletal protein/scaffold |
| Ubiquitin B-like | A0A8I5ZTA8\|A0A8I5ZTA8_RAT | Miscellaneous/unknown |
| Ubiquitin-40S ribosomal protein S27a | P62982\|RS27A_RAT | Miscellaneous/unknown |
| Ubiquitin-60S ribosomal protein L40 | A0A8I6G5G7\|A0A8I6G5G7_RAT | Miscellaneous/unknown |
| Uncharacterized protein | D3ZE08\|D3ZE08_RAT | Immunoglobulins |
| Uncharacterized protein | F1LTN6\|F1LTN6_RAT | Immunoglobulins |
| Uncharacterized protein | F1M229\|F1M229_RAT | Immunoglobulins |
| Uncharacterized protein | F1LQJ4\|F1LQJ4_RAT | Miscellaneous/unknown |
| Uncharacterized protein | A0A8L2UMW7\|A0A8L2UMW7_RAT | Miscellaneous/unknown |
| Uncharacterized protein | A0A8I6GC07\|A0A8I6GC07_RAT | Miscellaneous/unknown |
| Uncharacterized protein | A0A8I5ZW39\|A0A8I5ZW39_RAT | Immunoglobulins |
| Uncharacterized protein | A0A8I6AAU7\|A0A8I6AAU7_RAT | Metabolite interconversion enzyme |
| Uncharacterized protein | A0A8I6GKC8\|A0A8I6GKC8_RAT | Immunoglobulins |
| Uncharacterized protein | A0A8L2QLK6\|A0A8L2QLK6_RAT | Miscellaneous/unknown |
| Uncharacterized protein | A0A8I6GAC3\|A0A8I6GAC3_RAT | Metabolite interconversion enzyme |
| Vesicle amine transport 1-like | M0R3N4\|M0R3N4_RAT | Miscellaneous/unknown |
| Vimentin | P31000\|VIME_RAT | Cytoskeletal protein/scaffold |
| Vinculin | R9PXU6\|P85972\|VINC_RAT | Cytoskeletal protein/scaffold |
| Vitamin D-binding protein | Q68FY4\|P04276\|VTDB_RAT | Transfer/carrier protein/transporter |
| XPC complex subunit, DNA damage recognition and repair factor | D4A3D8\|D4A3D8_RAT | Miscellaneous/unknown |
| Zinc-alpha-2-glycoprotein | A0A8I5ZXF5\|F7F110\|F7F110_RAT | Defense/immunity protein |

**Supplementary Table S3:** The Top-20 most abundant proteins identified in native rat serum and CSF. RPA = relative protein abundance; Listed in alphabetical order.

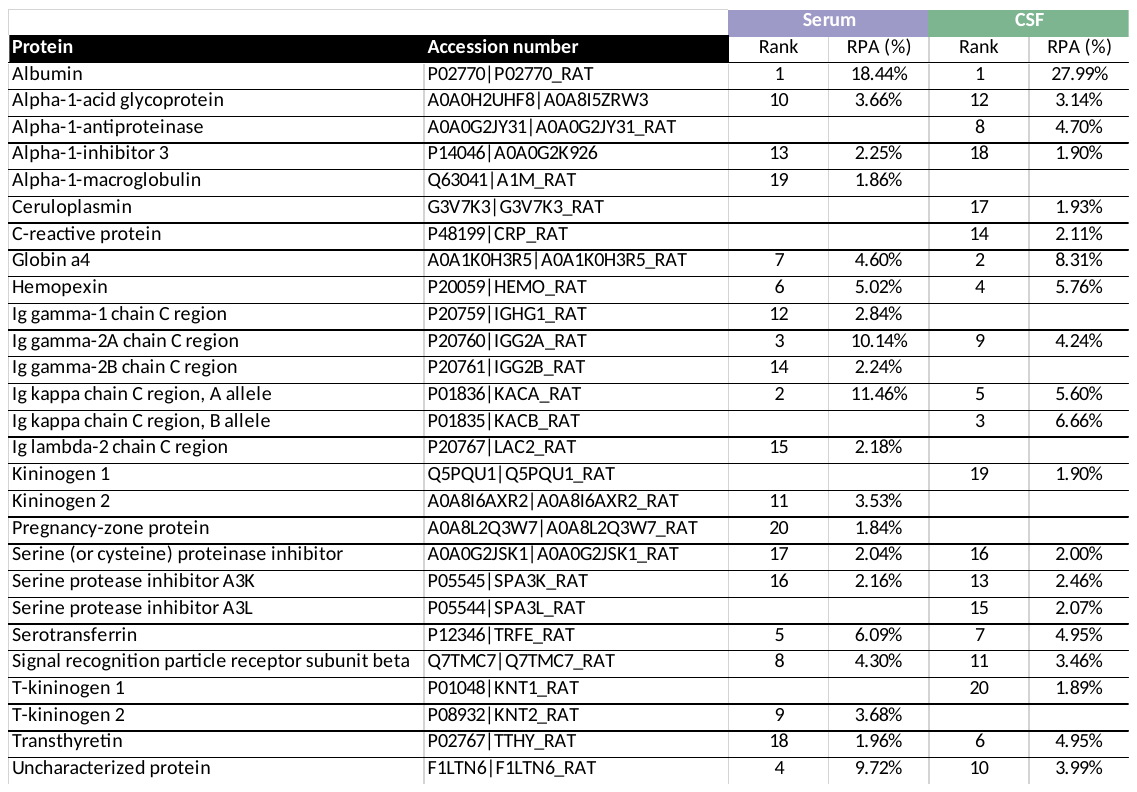

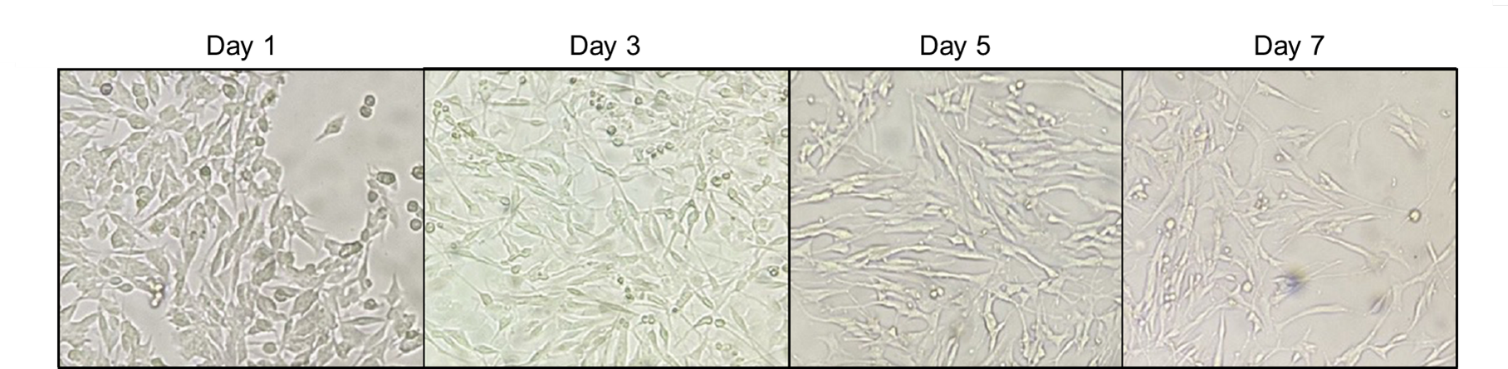

**Supplementary Figure S1:** Bright-field microscopy images showing the retinoic acid (RA)-induced differentiation of SH-SH5Y neuroblastoma cells into neurons. Over 7 days, RA-treated SH-SH5Y cells exhibit morphological changes, including long neurite outgrowth.

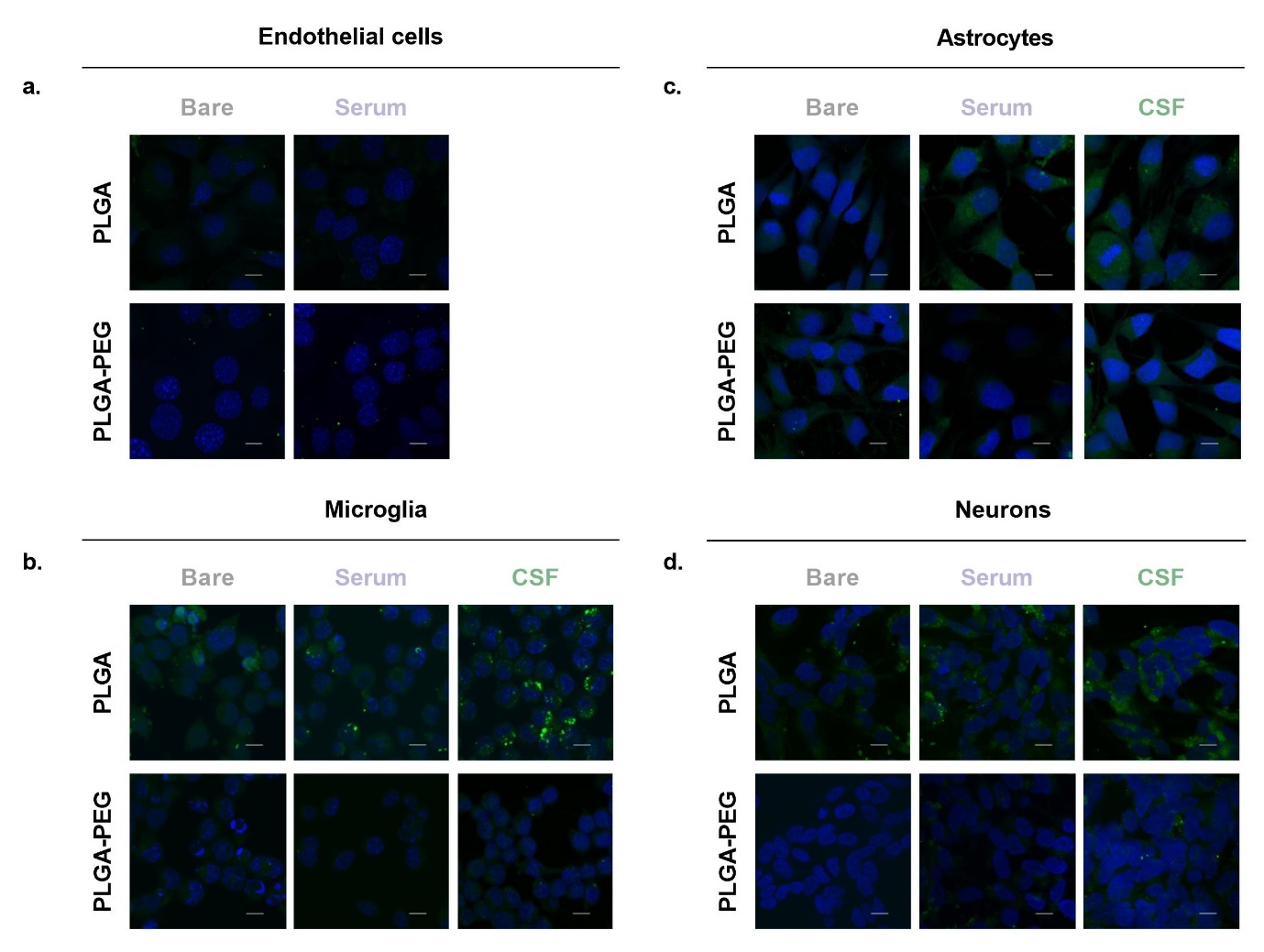
**Supplementary Figure S2:** Merged confocal fluorescence microscopy images showing the cellular uptake of FITC-loaded PLGA or PLGA-PEG nanoparticles (with and without coronas) by different brain cell types: **(a)** bEnd.3 brain endothelial cells; **(b)** BV2 microglia; **(c)**1321N1 astrocytes; and **(d)** SH-SY5Y-derived neurons. Nanoparticles (green); DAPI-stained cell nuclei (blue); Scale bars = 10 μm.

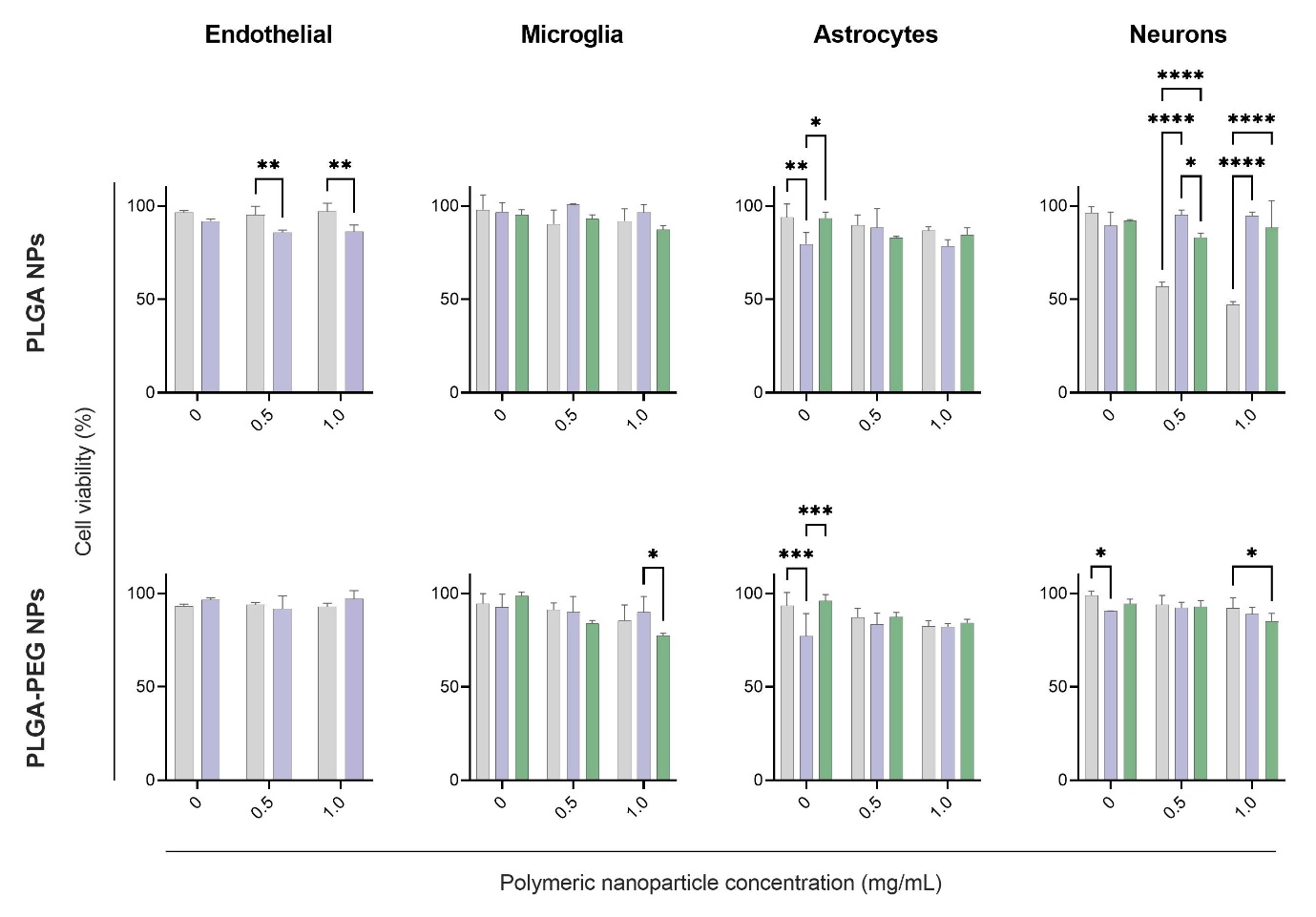

**Supplementary Figure S3:** Viability of different brain cell types determined by MTT assay. Cells were incubated with polymeric nanoparticles (with and without coronas) at varying concentrations (0, 0.5, and 1 mg/mL) for 24 h. Values represent means ± SD from three independent experiments. Error bars represent mean ± standard deviation, significance determined *via* 2way ANOVA with Šídák (Endothelial) or Tukey’s post-hoc analysis (*p<0.05, **p<0.01, ***p<0.001, ****p<0.0001).
